# Dysregulation of cellular iron predisposes chemotherapy resistant cancer cells to ferroptosis

**DOI:** 10.1101/2024.12.05.626685

**Authors:** Luke V. Loftus, Louis T.A. Rolle, Christopher Cherry, Michael Patatanian, Linda Orzolek, Bowen Wang, Kenneth J. Pienta, Sarah R. Amend

**Affiliations:** Cellular and Molecular Medicine Program, Johns Hopkins School of Medicine, Baltimore, MD; Cancer Ecology Center at the Brady Urological Institute, Johns Hopkins School of Medicine, Baltimore, MD; C M Cherry Consulting, Baltimore, MD; OMAPiX, Frederick, MD (Formerly Johns Hopkins University, Baltimore, MD & Psomagen, Rockville, MD); Pathobiology Program, Johns Hopkins School of Medicine, Baltimore, MD

## Abstract

Despite centuries of research, metastatic cancer remains incurable due to resistance against all conventional cancer therapeutics. Alternative strategies leveraging non-proliferative vulnerabilities in cancer are required to overcome cancer recurrence. Ferroptosis is an iron dependent cell death pathway that has shown promising pre-clinical activity in several contexts of therapeutic resistant cancer. However, ferroptosis sensitivity is highly variable across tissue types and cell state posing a challenge for clinical translation.

We describe a convergent phenotype induced by chemotherapy where cells surviving chemotherapy have similar transcriptomic signatures and dysregulated iron homeostasis, regardless of initial cell type or chemotherapy used. Elevated labile iron levels are counteracted by NRF2 signaling that does not alleviate the amount of labile iron. Selectively inhibiting GPX4 leads to uniform susceptibility to ferroptosis in surviving cells, highlighting the common reliance on lipid peroxidation defenses. Cellular iron dysregulation is a vulnerability of chemoresistant cancer cells that can be leveraged by triggering ferroptosis.

**STATEMENT OF SIGNIFICANCE:** We show that cells surviving chemotherapy are uniformly sensitive to ferroptosis, despite different sensitivity in untreated cells. Ferroptosis sensitivity surprisingly occurs alongside NRF2 signaling, likely due to loss of PCBP1 and accumulated labile iron. Labile iron accumulation as a driver of ferroptosis sensitivity constitutes another starting point for translation of ferroptosis.

## INTRODUCTION

Once cancer has metastasized, therapeutic resistance is the primary factor preventing durable cancer clearance. Nearly all current therapeutic strategies leverage the high proliferative rate of cancer cells to elicit apoptosis triggered by engagement of cell cycle checkpoints^1^. While anti-cancer therapies initially control tumor burden by eliminating the highly proliferative cancer cells, inevitably, a population of cancer cells evade apoptosis to survive, resulting in tumor recurrence. Non-apoptotic cell death pathways are cell cycle agnostic and are beginning to be explored to overcome therapeutic resistance in cancer.

Ferroptosis was officially named in 2012 as an iron-dependent form of cell death defined by unconstrained lipid peroxidation culminating in physical breakdown of plasma membrane integrity^2,3^. Iron reactivity is central to ferroptosis as iron both initiates lipid peroxidation and is required for amplification of lipid peroxidation through generation of alkoxyl radicals^4^. Glutathione peroxidase 4 (GPX4), the primary cellular defense against lipid peroxidation, prevents lipid hydroperoxide accumulation by converting to lipid alcohols^3^. Ferroptosis has gained considerable interest as an anti-cancer approach in a variety of drug tolerant cancer subpopulations all of which are heavily reliant on lipid peroxidation defenses^5–8^. Due to its metabolic underpinnings, however, mechanisms underlying ferroptosis sensitivity are highly variable across anatomical location, genetic background, and cellular phenotype over time^9–11^.

Iron utilization is as evolutionarily ancient as life itself. Planetary iron composition at the time of Earth’s formation was likely deterministic in the origin of biologic chemistry, and iron utilization is evident in the earliest unicellular lifeforms^12–15^. Subsequent global changes in iron bioavailability shaped selective pressures for iron acquisition, detoxification, and utilization that spurred evolution of symbiotic relationships and, ultimately, complex multicellular life on Earth^15^. That iron has retained its essential role throughout geological time and in nearly all species (save for two bacterial life forms^16^) highlights its irreplaceable role in numerous vital functions including DNA replication, transcription, redox sensing, and mitochondrial metabolism^17–19^. However, iron’s unique electron transfer properties that enable life also indiscriminately generate toxic radicals when iron is freely reactive^20,21^. Thus, throughout the biochemical and geochemical coevolution of iron utilization, cellular programs to manage irons access to oxygen and oxidation state are critical to avoid toxicity.

In humans, cellular iron is imported primarily through endocytosis of the transferrin receptor and liberation of iron in mature lysosomes. Minimal amounts of ferrous iron exist in a dynamic and exchangeable labile iron pool (LIP) before incorporation into cofactors as well as directly into protein active sites^18^. Even in the LIP, iron is predominantly bound to glutathione to minimize spurious reactivity^22,23^. When free iron accumulates intracellularly, it is detoxified and stored as Fe^3+^ in ferritin proteins that evolved along the ancient history of iron utilization^24^. Poly(rC)-binding protein 1 (PCBP1) is the primary iron chaperone shown to directly bind and deliver Fe-GSH to ferritin and several apoenzymes^23,25,26^.

Cancer has long been appreciated as iron addicted to support proliferation and increasing biomass. Less is known about intracellular management of iron during cancer progression as well as how cells handle this vital, but potentially toxic, resource when reacting to therapeutic stress. We have previously shown that cells surviving single dose chemotherapy uncouple DNA replication and cell division leading to increased ploidy and cell size and the surviving cells are resistant to subsequent chemotherapy^27^. To identify convergent phenotypes and shared vulnerabilities of chemotherapy resistance, we evaluated two cell lines of different tumor types (PC3; metastatic prostate cancer cell line and MDA-MB-231; metastatic breast cancer cell line) and two classes of chemotherapy (alkylating agent cisplatin and microtubule stabilizer docetaxel). We found that cells surviving chemotherapy have high amounts of labile iron regardless of cell line or chemotherapy treatment. We performed single cell RNA sequencing of these populations and identified canonical cellular responses to iron toxicity including NRF2 signaling, upregulation of glutathione (GSH) biosynthesis and ferritin expression in all settings. Despite the physiological response to iron accumulation in surviving cells, labile iron was not attenuated, likely due to a relative loss of the PCBP1 chaperone. Labile iron burden in resistant cells conferred uniform sensitivity to ferroptosis via GPX4 inhibition, regardless of originating cell line or chemotherapy. This study highlights labile iron accumulation as a driver of ferroptosis sensitivity in chemoresistant cells, despite active NRF2 signaling. For ferroptosis to translate into a viable clinical option in metastatic cancer settings, timing of intervention will be critical to maximize efficacy and enable treatment of multiple tumor types.

## RESULTS

### Cells surviving chemotherapy converge on a similar transcriptomic signature

PC3 prostate and MDA-MB-231 breast cancer cell lines were treated with cisplatin or docetaxel for 72 hours, then drug was removed, and cells were followed up to 10 days post-treatment (DPT). Consistent with previous reports^27,28^, surviving cells increased in size with time (Supplementary Fig. S1). Single cell RNA sequencing was performed at one, five, and ten DPT via combinatorial barcoding to accommodate cell size. We then used unsupervised dimensionality reduction to identify major transcriptomic signatures and plotted clusters using Uniform Manifold Approximation and Projection (UMAP). To shed light on biological processes enriched in cells that survive treatment, we performed gene set enrichment analysis (GSEA), cell cycle scoring, and Ucell scoring^29,30^. While MDA-MB-231 control cells clustered separately (in clusters 1 and 6), cells surviving chemotherapy did not cluster solely by timepoint indicating a heterogenous, dynamic response to chemotherapy (Figure 1A and 1B). Cells 1 and 5 days post cisplatin resided in multiple clusters defined by cell cycle signatures similar to untreated cells, enriched epithelial-mesenchymal transition, oxidative phosphorylation, and a minority cluster defined by unfolded protein response genes (Fig. 1B and 1C, Supplementary Fig. S2). Docetaxel treated cells at 1 DPT primarily resided in one cluster with decreased cell cycle genes and elevated expression of ribosome and spliceosome genes (Fig. 1B, Supplementary Fig. S2). By 10 DPT of either chemotherapy, nearly all surviving MDA-MB-231 cells cluster together (clusters 0 and 2), with one distinct 10 day post cisplatin cluster (cluster 4).

**Figure 1.**
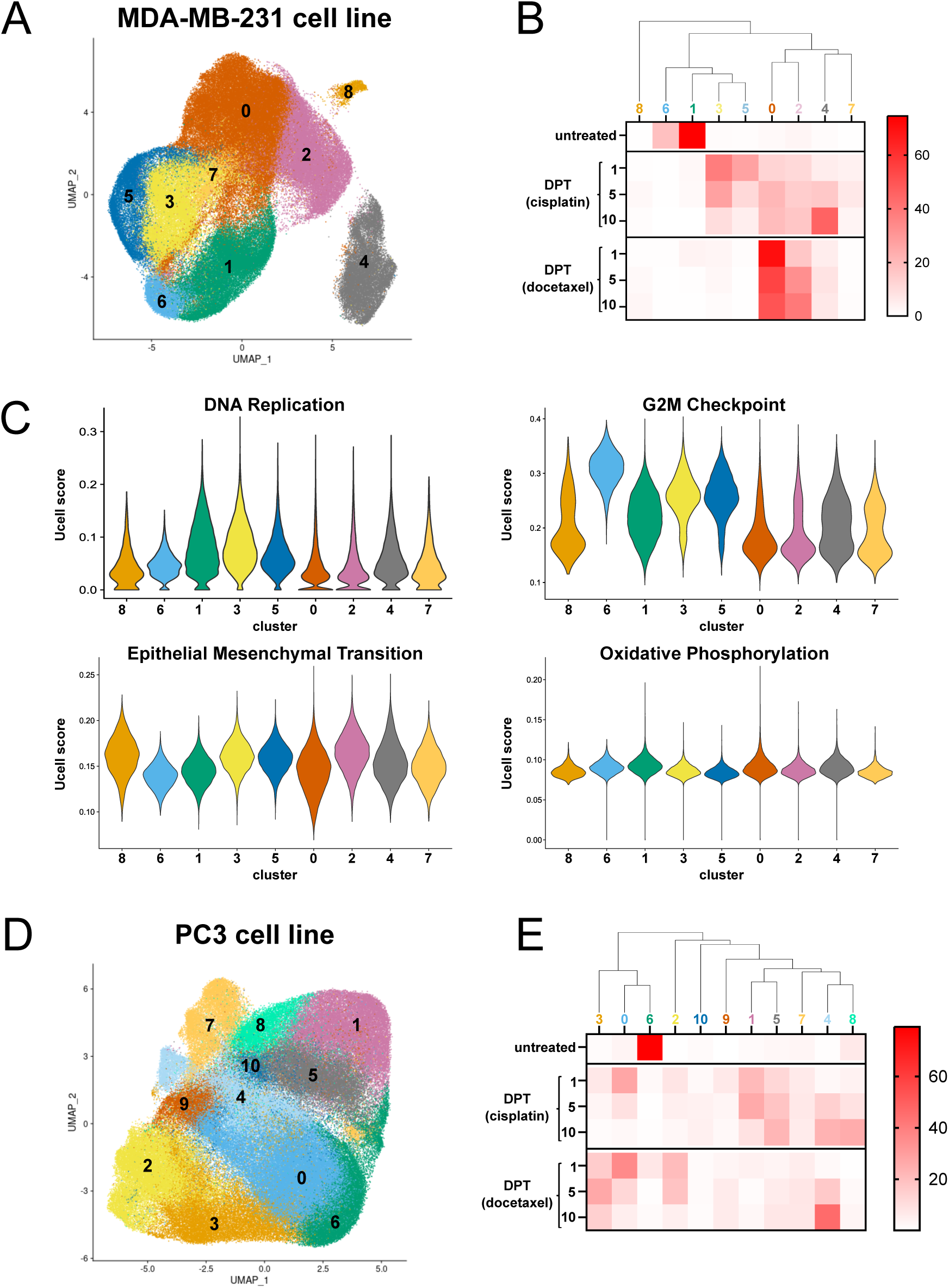
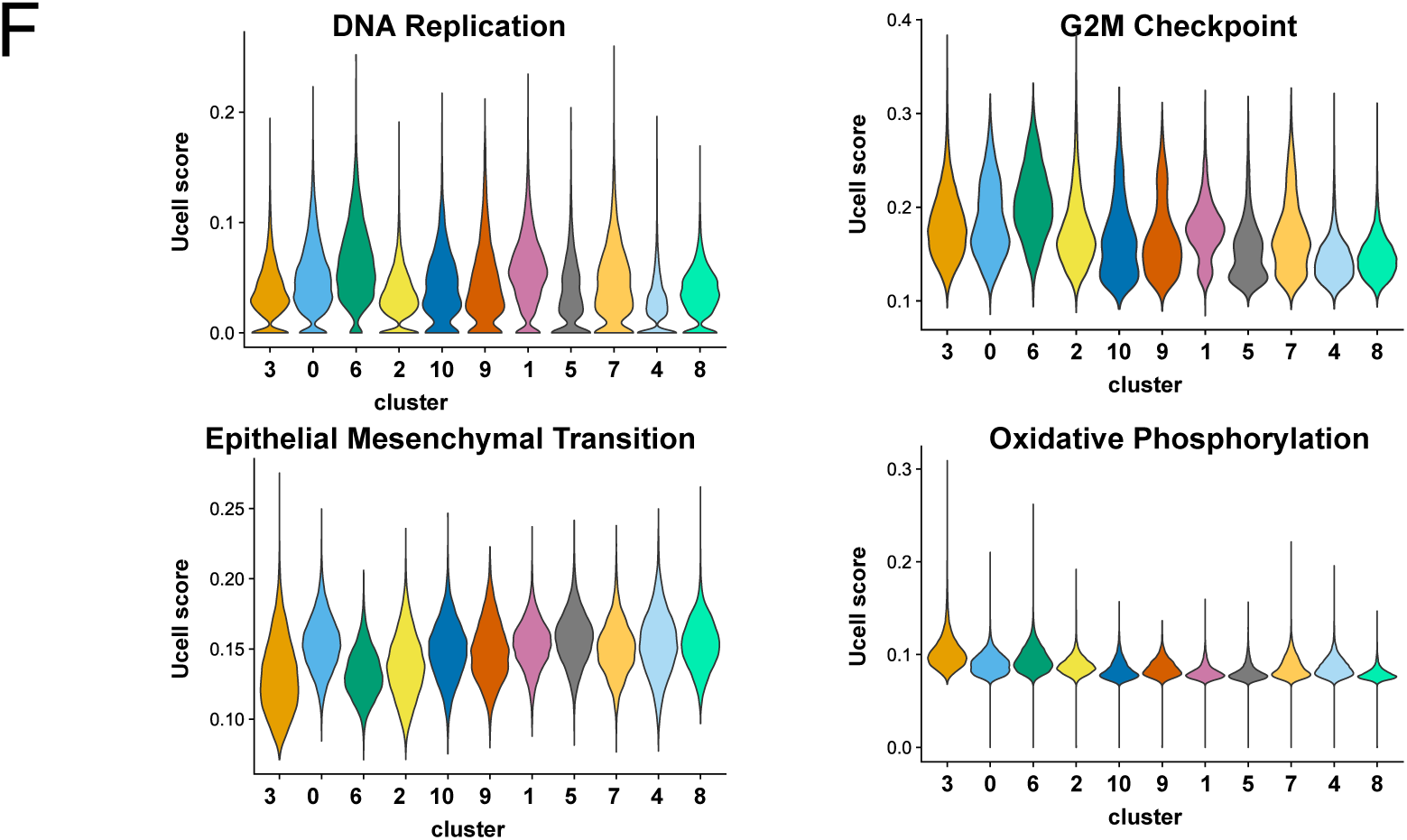
Cells surviving chemotherapy have diverse transcriptomic signautres not confined to treatment or timepoint alone. **A.** scRNAseq UMAP projection for MDA-MB-231 cells before and after cisplatin or docetaxel treatment. **B.** Distribution of samples across clusters in A. Heatmap coloring represents percent of sample in each cluster. **C.** Ucell score of selected genesets across each cluster in A. **D.** scRNAseq UMAP projection for PC3 cells before and after cisplatin or docetaxel treatment. **E.** Distribution of samples across clusters in A. Heatmap coloring represents percent of sample in each cluster. **F.** Ucell score of selected genesets across each cluster in A.

Like in MDA-MB-231 cells, untreated PC3 cells clustered independently (cluster 6). Cells 1 day post cisplatin occupied several clusters while docetaxel treated cells had relatively restricted clustering (Fig. 1D and 1E). A single cluster with notable overlap of cells 1 DPT was also present, with enriched oxidative phosphorylation, as well as decreased cell cycle signatures relative to untreated cells (Fig. 1E and 1F, Supplementary Fig. S3). Also similar to MDA-MB- 231 surviving cells, PC3 cells 10 DPT overlapped in clustering, with cisplatin 10 DPT cells residing in one additional cluster (cluster 8).

While cell line specific and chemotherapy specific differences were evident as expected, the overlapping gene signatures at 10 DPT was most interesting as it suggests a shared phenotype of surviving cells. Comparing GSEA of cells 10 DPT versus their respective untreated cell line, we found 15 gene sets that were common to all 10 DPT surviving cells, regardless of cell line or treatment types. These shared gene sets comprise decreased cell cycle genesets, including Myc Targets and ribosome components, and enriched inflammatory genesets (Fig. 2A, Supplementary Table T1). We used Ucell scoring with a refined geneset list to visualize changes in gene expression over cell survival timepoints (Fig. 2B). These transcriptomic signatures arise over time and have less variance by 10 DPT, reflecting a convergent phenotype regardless of tumor type or chemotherapy (Fig. 2B, Supplementary Fig. S4). A refined geneset list also identified enrichment in autophagy genes as part of this convergent phenotype.

**Figure 2.**
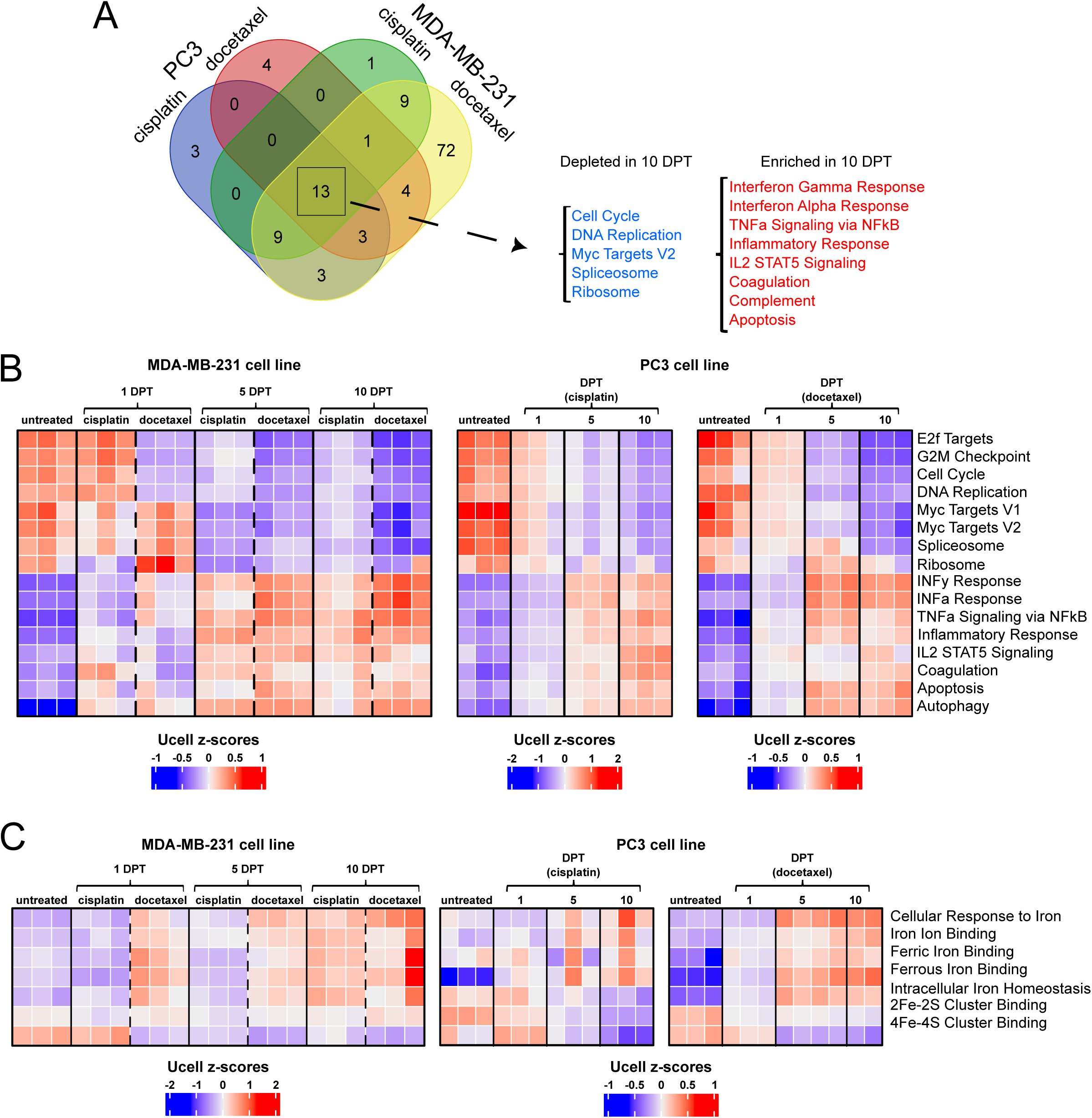
Surviving cells 10 DPT converge on similar transcriptomic signatures. **A.** scRNAseq GSEA analysis of 10 DPT cells compared to their respective untreated cells. Numbers indicate the number of significant genesets. **B.** Ucell analysis for overlapping genesets from A. plus autophagy geneset. Ucell mean was calculated per sample and displayed as mean value z-scored across samples. Each column is a biological replicate. **C.** Ucell analysis for iron handling genesets. Ucell mean was calculated per sample and displayed as mean value z-scored across samples. Each column is a biological replicate.

Active transcription with numerous enriched genes, co-existing with enriched autophagy and decreased ribosome signatures, in cells 10 DPT reflects a decrease in protein translation relative to transcription that is commonly observed in cells with increased genomic content^31^. An imbalance between transcriptome and proteome also creates imbalances in protein cofactors including iron which is essential for a multitude of proteins across the proteome^17^. We evaluated iron handling genesets and found changes in intracellular iron homeostasis and iron binding (Fig. 2C). Transferrin receptor is downregulated in cells 10 DPT, but some contexts have increased CD44, an alternate route for iron import (Supplementary Fig. S5). Iron transporters DMT1 (SLC11A2) and to a lesser extent Lipocalin 2 (LCN2) are enriched at 10 DPT in both cell lines and both treatment types, implying a high requirement for iron transport in surviving cells. HMOX1 (Heme Oxygenase 1, degrades heme and releases ferrous iron) is minimally expressed (Supplementary Fig. S5). SLC40A1 (Ferroportin, ferrous iron export) also is not appreciably expressed in surviving cells, consistent with conventional iron export being restricted to enterocytes, hepatocytes, and macrophages^18^.

### Cells surviving chemotherapy have high labile iron

Cells surviving chemotherapy require iron to increase cellular biomass but showed transcriptomic signatures for mismanagement of cellular iron (Fig. 2C, Supplementary Fig. 4C). Physiological regulation of iron maintains low levels of labile iron to accommodate protein synthesis demands but avoid spurious reactivity and oxidative damage^18^. We assessed labile iron by live cell imaging (Fig. 3) and an orthogonal lysate-based assay (Supplementary Fig. 6). Labile iron levels were higher in untreated PC3 than MDA-MB-231 cells and was modestly elevated at 1-DPT in both cell lines, regardless of chemotherapy treatment, consistent with de-compartmentalization of iron following acute stress^20^ (Fig. 3A-C, Supplementary Fig. S6A-B). Labile iron increased dramatically from 5 to 10 DPT, in both of cell lines and chemotherapy groups (Fig. 3A-C, Supplementary Fig. S6B), indicating a common mismanagement of cellular iron. Labile iron accumulates primarily in mitochondria and ER due to iron-containing proteins and protein assembly in both compartments^17^. At 10 DPT, labile iron had visibly punctate staining that exceeded ER boundaries, regardless of cell line or chemotherapy, further implying a dysregulation in iron homeostasis. (Fig. 3D, Supplementary Fig. S6C). Punctate labile iron staining in cells 10 DPT showed largely overlaps with active lysosomes (Figure 3D, Supplementary Fig. S6C).

**Figure 3.**
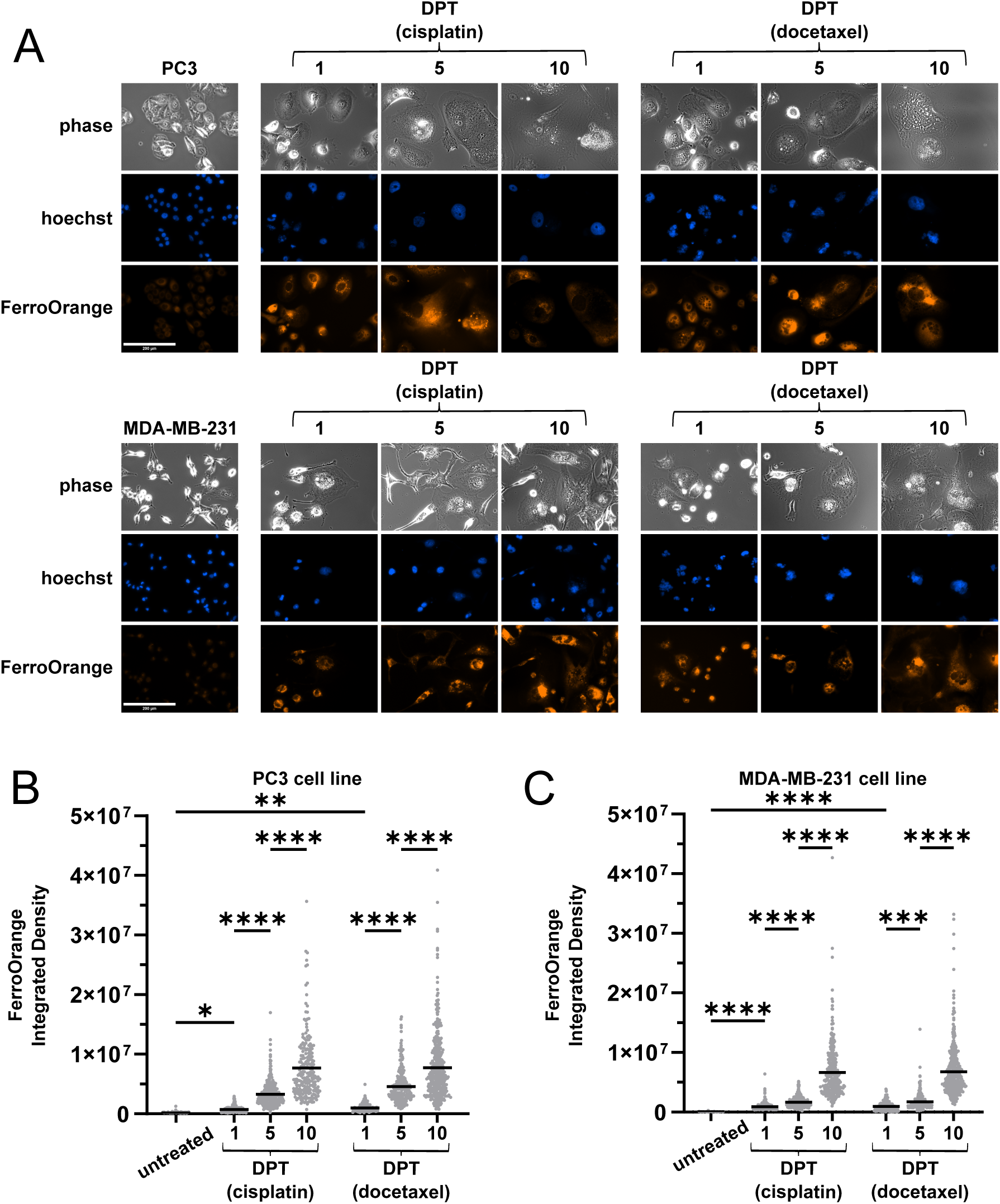
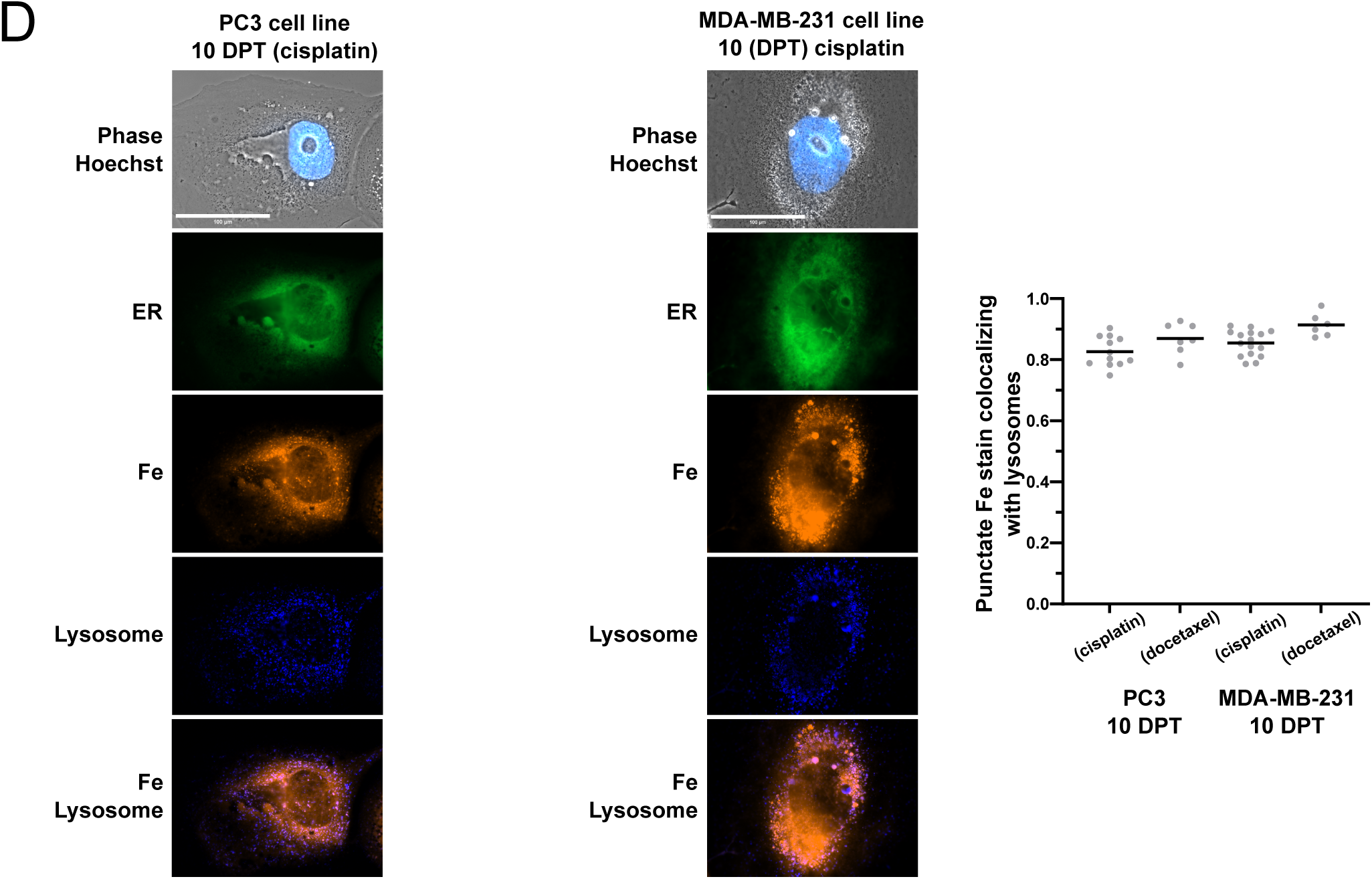
Cells surviving chemotherapy have high amounts of labile iron. **A.** Live cell images of PC3 and MDA-MB-231 cells before and after cisplatin or docetaxel using FerroOrange. **B, C.** Quantification of FerroOrange integrated density from A. in PC3 (**B**) or MDA-MB-231 (**C**) cells before and after chemotherapy. Data displays individual cells (dot) and mean (bar) from three biological replicates. **D.** Live cell images with FerroOrange (Fe), ERTracker Green (ER), and LysoTracker (Lysosome) of PC3 and MDA-MB-231 cells 10 days after cisplatin treatment. (right) Quantification of punctate FerroOrange staining colocalizing with LysoTracker staining. Data is Manders Colocalization Coefficient for fraction of punctate iron stain colocalized with lysosome.

### Ferritin expression does not attenuate labile iron levels

In response to labile iron accumulation, cells detoxify and store Fe^3+^ in ferritin^32^. Untreated cells lines and early DPT do not have appreciable ferritin expression, but by 10 DPT there is a marked increase in ferritin across all samples (Fig. 4A-B, Supplementary Fig. S6D). Paradoxically, this elevated ferritin expression does not accompany a decrease in the LIP in cells 10 DPT (Fig. 3A-C, Supplementary Fig. S6B). In addition, while Transferrin Receptor RNA is downregulated in surviving cells (Supplementary Fig. S5C), it is unchanged at the protein level (Supplementary Fig. S6E), suggesting a disconnect between surviving cells’ requirement for iron utilization vs storage.

**Figure 4.**
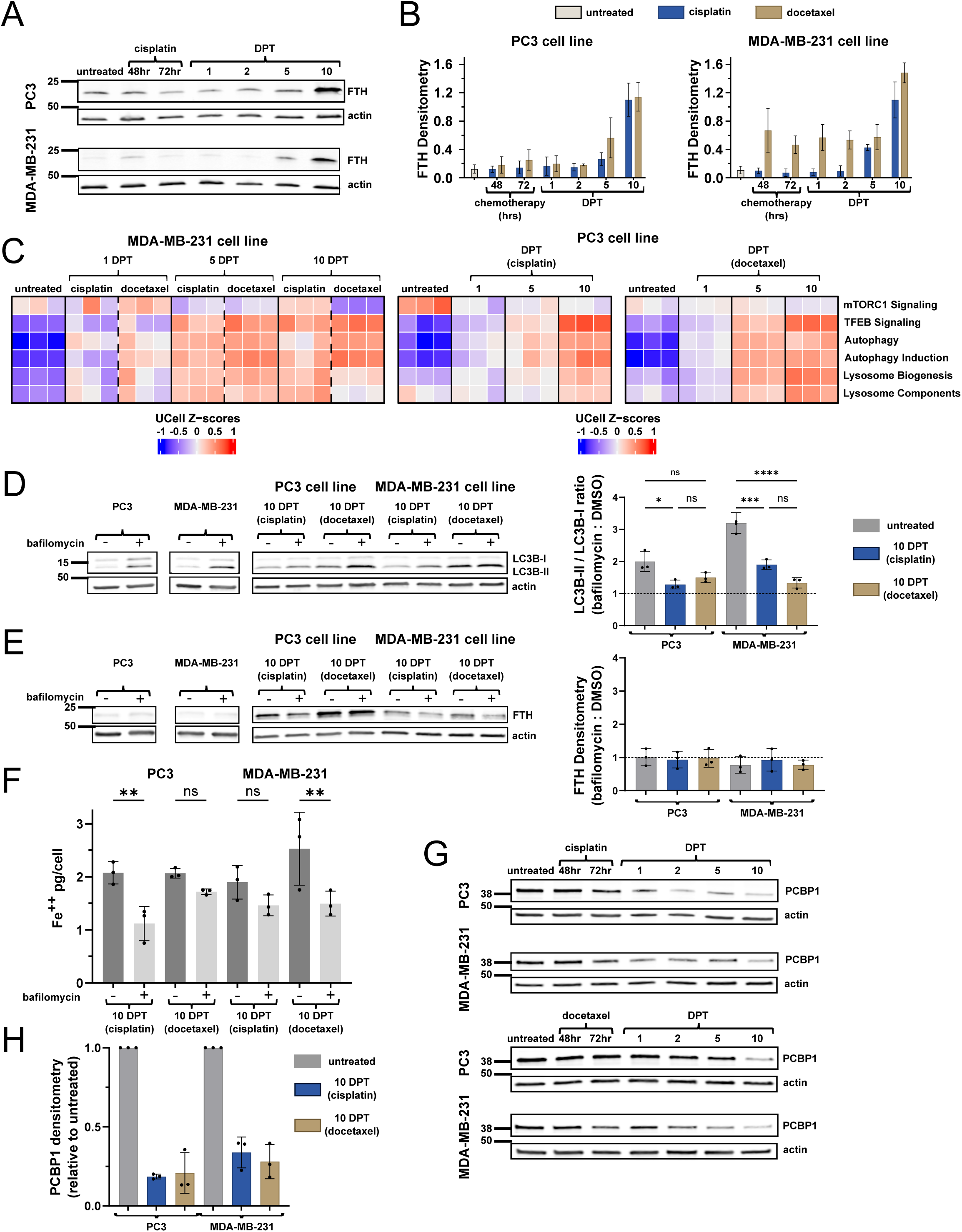
Cells surviving chemotherapy have high ferritin expression that is not being turned over by autophagy. **A.** Western blot for ferritin in PC3 and MDA-MB-231 cells before and after cisplatin. **B.** Quantification of ferritin expression from A. Data is mean and standard deviation of three biological replicates. **C.** scRNAseq analysis of autophagy related genesets. Ucell score for each geneset was calculated and represented as z-scored heatmap across samples. **D.** Western blot for LC3B expression with and without bafilomycin treatment in PC3 and MDA-MB-231 cells surviving chemotherapy. (right) Quantification of LC3B-II / LC3B-I ratio in bafilomycin / DMSO condition. Dotted line at 1 represents no change from DMSO treated LC3B ratio. **E.** Western blot for ferritin expression with and without bafilomycin treatment in PC3 and MDA-MB-231 cells surviving chemotherapy. (right) Quantification of ferritin in bafilomycin / DMSO condition. Dotted line at 1 represents no change from DMSO treated ferritin expression. **F.** Labile iron quantification with and without bafilomycin treatment in PC3 and MDA-MB-231 cells surviving chemotherapy. **G.** Western blot for PCBP1 in PC3 and MDA-MB-231 cells surviving chemotherapy. **H.** Quantification of PCBP1 expression from G. in 10 DPT cells relative to respective untreated cells.

To further investigate iron homeostasis, we next looked at autophagy as selective ferritin degradation (ferritinophagy) is a well characterized contributor to labile iron accumulation^3^. Initial transcriptomic characterization identified enriched autophagy gene expression in cells 10 DPT (Fig. 2B) and additional autophagy related genesets, including TFEB signaling and lysosomal biogenesis, were also enriched (Fig. 4C). Cells 10 DPT have an elevated LC3B-II to LC3B-I ratio indicating increased autophagosome formation (Supplementary Fig. S7B). However, upon inhibition of autophagy with bafilomycin, there is less accumulation of LC3B at 10 DPT than in untreated cells, indicative of decreased autophagic flux (Fig. 4D). In all groups, bafilomycin treatment does not lead to accumulation of ferritin, meaning ferritin is not being degraded through autophagy (Fig. 4E). Accordingly, autophagy is not contributing to the elevated labile iron levels in surviving cells either (Fig. 4F). Bafilomycin interferes with lysosomal biology, not only autolysosomes, which, alongside the lysosomal accumulation of iron, likely explains partial labile iron modulation by bafilomycin in 10 DPT surviving cells (Fig. 4F).

In the absence of ferritinophagy and other known drivers of iron accumulation (Supplementary Fig. S5), we evaluated iron trafficking as a mechanism underlying increased iron in surviving cells. PCBP1 is the primary iron chaperone known to deliver Fe-GSH complexes to ferritin^25^. PCBP1 is required for iron loading into ferritin and loss of PCBP1 causes poor ferritin iron loading, increased labile iron pool and lipid peroxidation^25,33,34^. PCBP1 is ubiquitously expressed at high levels in mammalian cells as is necessary for iron chaperone activity^23^, and indeed PCBP1 protein is abundant in untreated cancer cell lines (Fig. 4G). However, PCBP1 expression waned over time following cisplatin or docetaxel treatment, with cells 10 DPT being significantly depleted of PCBP1 across all contexts (Fig. 4G-H). PCBP1 is not transcriptionally downregulated nor is it being turned over by autophagy (Supplementary Fig. S8).

Taken together, these data suggest that while cells surviving chemotherapy are responding to labile iron accumulation, they are not effectively sequestering iron, likely due to a relative loss in PCBP1 chaperone.

### Surviving cells have increased NRF2-mediated antioxidant response

Given excessive labile iron in cells that survive chemotherapy, we next investigated cellular mechanisms to mitigate iron toxicity. Transcription factor NRF2 is a master coordinator of the cellular antioxidant response to oxidative damage, including from heavy metal toxicities^35,36^. MDA-MB-231 cells have slightly higher NRF2 expression than PC3 cells both in untreated cells and during chemotherapy and 1 DPT timepoints (Fig. 5A, Supplementary Fig. S9A). By 10 DPT, surviving cells have high NRF2 expression regardless of cell line or chemotherapy (Fig. 5A, Supplementary Fig. S9A). Deferoxamine (DFO) is an iron chelator that attenuates cellular iron import through endocytosis and chelation of iron released from transferrin in nascent lysosomes^3^. DFO treatment for six hours partly attenuated NRF2 expression in PC3 cells 10 DPT, but not MDA-MB-231 cells 10 DPT, indicating potential differences in active iron import contributing to NRF2 activity (Fig. 5B). Consistent with high NRF2 expression, known transcriptional targets of NRF2^37^ are enriched in surviving cells 10 DPT regardless of cell line or chemotherapy (Fig. 5C, Supplementary Fig. S9B-C). Transcriptionally enriched NRF2 targets include Ferritin (FTH), GSH biosynthesis enzymes including catalytic and regulatory subunits of Glutamate-cysteine ligase (GCLC, GCLM), and p62 (SQSTM1) for coordinating damaged protein clearance (Fig. 5C, Supplementary Fig. S9B- C). Ferritin and p62 are likewise highly expressed at the protein level (Fig 4A-B, Supplementary Fig. S6D and S9D), suggesting a connection between oxidative stress, inefficient autophagy, and protein clearance. In MDA-MB-231 cells treated with docetaxel, cells 1 DPT already have strong enrichment of NRF2 targets, but overall fewer cells express each target gene; by 10 DPT, most NRF2 target genes are still enriched (Supplementary Fig. S5B).

**Figure 5.**
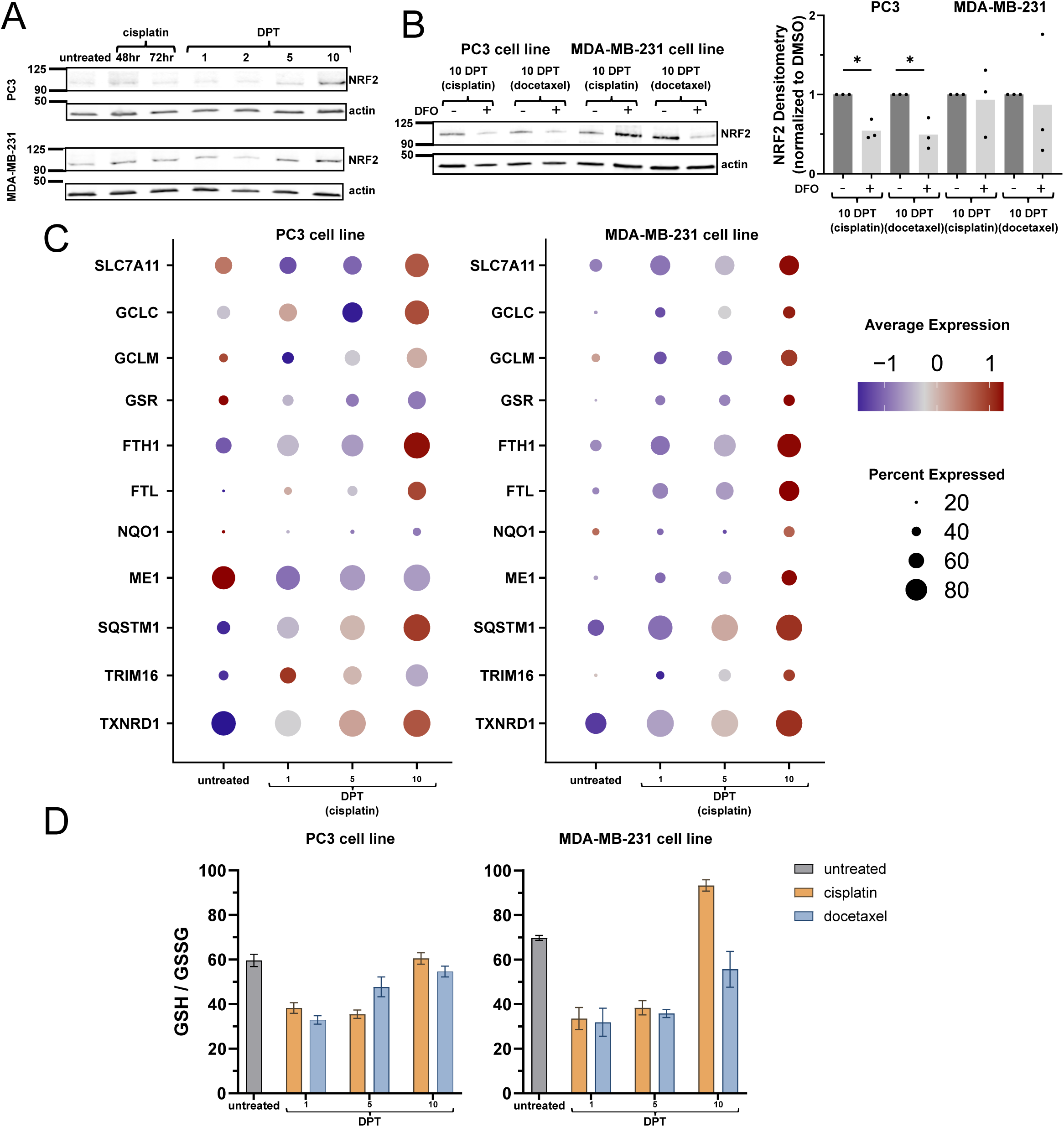
10 DPT surviving cells have high NRF2 expression and enriched NRF2 transcriptional targets. **A.** Western blot for NRF2 in PC3 and MDA-MB-231 cells before and after cisplatin. **B.** Western blot for NRF2 following 6 hours of deferoxamine mesylate (DFO) treatment in cells 10 days post-chemotherapy. **C.** NRF2 target gene expression in PC3 or MDA-MB-231 cells surviving cisplatin. Each gene represented by z-scored distribution of average normalized expression across samples (color) and percentage of cells with detected transcript (size). **D.** GSH : GSSG ratio in PC3 and MDA-MB-231 cells following cisplatin or docetaxel treatment.

### Oxidative stress in surviving cells

The ratio of reduced to oxidized glutathione (GSH) is a widely used indicator of oxidative stress. We measured GSH ratio by plate based assay and found that cells 1 and 5 DPT have a compromised GSH ratio, indicative of general oxidative stress from chemotherapy, but by 10 DPT the GSH ratio is restored to similar levels as untreated cells (Fig. 5D). Restoration of the GSH ratio is consistent with the transcriptional enrichment of GSH biosynthesis pathway at 10 DPT (Fig. 5C, Supplementary Fig. S9B-C), including MDA-MB-231 cells 10 days post docetaxel having a lower restoration of GSH ratio. Reduced GSH is the most abundant antioxidant in cells with well-known roles in direct detoxification and regeneration of antioxidant enzyme active sites. GSH is also the primary ligand for ferrous labile iron serving as a critical non-chelating ligand that enables intracellular iron trafficking and selectivity over manganese for enzyme incorporation^22,38,39^. It is hypothesized GSH’s primary role may be for iron handling, secondary to a redox buffer, with the high cytosolic abundance preserve its iron handling functions even during fluctuations from redox stressors^40^. Based on these results, we hypothesize that in surviving cells 10 DPT, reduced GSH is both occupied by the excessive amount of labile iron, particularly with the relative loss of PCBP1, and serving its traditionally described antioxidant roles to counteract oxidative damage from accrued DNA damage and iron catalyzed reactions.

### Cells surviving chemotherapy are vulnerable to ferroptosis

Labile iron reactivity is connected to lipid peroxidation through Fenton reactions creating free radicals, catalyzing enzyme-dependent lipid peroxide formation, and direct reaction with lipid peroxides to generate alkoxyl radicals^41^. Given high labile iron and reduced GSH pools in cells surviving chemotherapy, we hypothesized that cell survival is reliant on lipid peroxidation defense systems. We assessed lipid peroxidation using C11 BODIPY reagent and found increased lipid peroxidation in cells 10 DPT (Fig. 6A). Thus, despite the accumulated labile iron, cells 10 DPT are effectively counteracting lipid peroxidation prior to any intervention.

**Figure 6.**
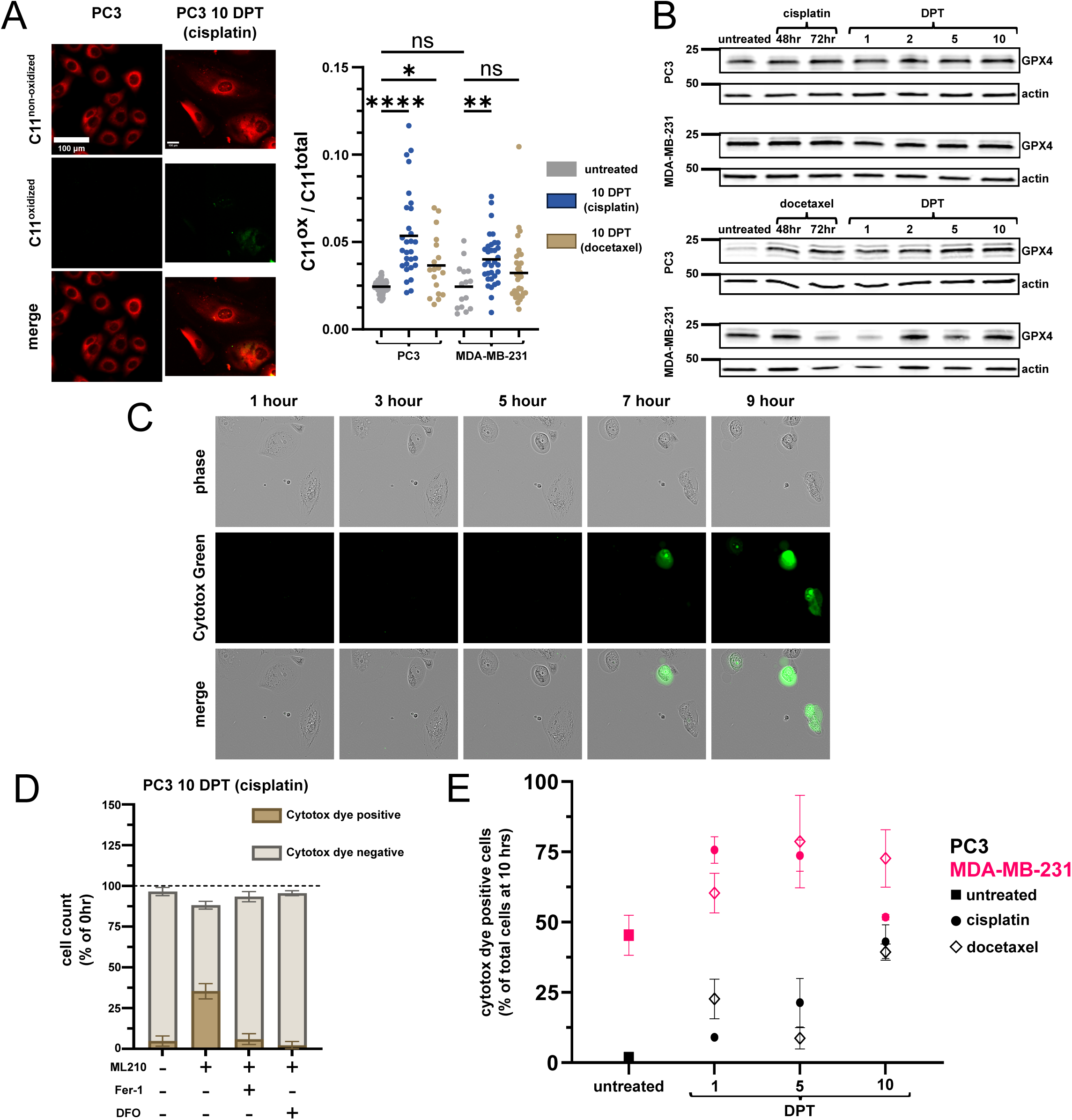
Surviving cells 10 days post chemotherapy are susceptible to ferroptosis regardless of cell line of origin. **A.** C11 Bodipy imaging in PC3 cells untreated and 10 days post cisplatin. Quantification is ratio of oxidized (green) to total (green + red) C11 fluorescence. **B.** GPX4 expression in PC3 or MDA-MB-231 cells surviving cisplatin or docetaxel. **C.** Timelapse images of PC3 cells 10 days post cisplatin treated with ML210. Cytotox dye incorporation indicates membrane permeability. **D.** Quantification of live cell imaging for PC3 cells 10 days post cisplatin treated with, from left to right, DMSO (0.1%); ML210 (1uM); ML210 and Fer-1 (1uM); ML210 and DFO (50uM). Fer-1 = Ferrostatin-1. DFO = Deferoxamine. **E**. Percent of Cytotox dye positive cells following ML210 (1uM) treatment across all samples.

GPX4 is the only mammalian antioxidant enzyme capable of cleansing lipid peroxidation. GPX4 expression is largely unchanged across all cell lines and timepoints (Fig. 6B, Supplementary Fig. S10A), suggesting lipid peroxidation is combatted through regeneration, rather than increased expression, of GPX4. We utilized the GPX4 inhibitor ML210 to assess ferroptosis vulnerability. ML210 is a masked nitrile-oxide electrophile that is converted intracellularly to a transient active compound which then covalently modifies the GPX4 active site^42,43^. Cells were treated with 1uM dose of ML210 and ferroptosis was quantified over a 10- hour period. As expected, GPX4 inhibition causes an increase in lipid hydroperoxide abundance by 6 hrs (Supplementary Fig. S10B). Ferroptosis was quantified by time lapse imaging to account for cell loss from apoptosis (especially in early recovery timepoints following chemotherapy) and visualize loss of membrane integrity. Susceptible cells undergo membrane blowouts and full loss of membrane integrity, as detected by uptake of a cell impermeable dye, and remain adherent at the 10-hour timepoint (Fig. 6C, Supplementary Fig. S10C, Supplementary Movie S1-4). To confirm ferroptosis as the mechanism of cell death, cell death was fully prevented in the presence of Ferrostatin-1 (lipid alkoxyl radical scavenger)^44^ or deferoxamine (iron chelator) in all samples (Fig. 6D, Supplementary Fig. S11, Supplementary Movie S5-6).

Consistent with the known ferroptosis sensitivity of the MDA-MB-231 cell line^45,46^, we find 45% of untreated MDA-MB-231 cells die due to ferroptosis with GPX4 inhibition (Fig. 6E). Chemotherapy increased sensitivity to ferroptosis with a peak sensitivity at 5 DPT where around 75% of MDA-MB-231 cells undergo ferroptosis regardless of chemotherapy group. At 10 DPT, cisplatin treated cells were less vulnerable to ferroptosis, at similar levels to untreated MDA-MB-231 cells. Docetaxel treated cells retained much of their susceptibility likely due to the (relatively) lower GSH ratio and marked transcriptional differences from cisplatin treated cells 10 DPT (Fig. 2A, Fig. 6E).

Untreated PC3 cells are agnostic to GPX4 inhibition and cisplatin only modestly increased sensitivity at 1 and 5 DPT. 23% of PC3 cells 1 days post docetaxel died to ferroptosis, but by 5 days post docetaxel sensitivity was only 8%. PC3 cells 5 DPT docetaxel were not significantly more sensitive to ferroptosis than untreated PC3 cells, likely due to ferritin, GSH pathway, and GSH ratio enrichment at 5 DPT in this context (Fig. 6E). By 10 DPT, regardless of chemotherapy, surviving cells were more vulnerable to ferroptosis at similar levels to MDA-MB- 231 cells 10 days post cisplatin (Fig. 6E).

Similar vulnerability to ferroptosis at 10 DPT in both cell lines and both treatment groups, following different dynamics at 1 and 5 DPT, aligns with the labile iron driven reliance on GPX4. Critically these cells are not spontaneously dying from ferroptosis (Fig. S11 DMSO treated conditions), reinforcing the fact that surviving cell populations are managing their labile iron burden by relying on GPX4 and its regeneration by GSH. Ferroptosis has been reported as a form of immunogenic cell death in some settings that, upon loss of membrane integrity, may increase immunogenicity and activate initiate immunity^47^. We evaluated common molecular markers of immunogenicity including ATP and HMGB1 release, as well as M-CSF and IL-2, in cells 10 DPT co-cultured with PBMCs both before and after ferroptosis (Supplementary Fig. S12). Ferroptotic cell death at 10 DPT did not release commonly recognized immunogenic molecules, although more research is warranted to identify additional immunogenic aspects of ferroptosis.

## DISCUSSION

In this study, we sought to leverage ferroptosis as a proliferation- and apoptosis-agnostic mechanism for elimination of resistant cancer. We utilized two different cell lines and chemotherapies and identified a convergent phenotype that arises over time, such that cells that have survived 10 days post chemotherapy, regardless of cell line or chemotherapy, display similar morphology and transcriptomic signatures. Surviving cells 10 DPT have high amounts of labile iron and display characteristics of iron toxicity, where NRF2 activation leads to GSH pathway enrichment and high ferritin expression, yet ferritin is not alleviating labile iron levels. Nor can labile iron levels be explained by well known drivers of labile iron accumulation such as HMOX1 and ferritinophagy. Instead, we find that a relative loss of PCBP1 expression is the most likely cause of labile iron accumulation. A shared labile iron burden in this convergent phenotype creates a similar reliance on GPX4, and thus vulnerability to ferroptosis, despite different sensitivities to ferroptosis in untreated cell lines. Our study highlights a convergent phenotype with a shared, labile iron driven, vulnerability to ferroptosis. Ferroptosis sensitivity in this setting cannot be explained by EMT driven modulation of membrane fluidity or a downregulation of GSH pathways (Supplementary Fig. S13), as is seen in other contexts of resistant cancer sensitivity to ferroptosis^5–7,46,48^. Instead, vulnerability to ferroptosis counterintuitively occurs along with NRF2 activation, high GSH levels and high ferritin expression in surviving cells.

PCBP1 is the primary cellular iron chaperone and is required for iron loading into ferritin, cytosolic [2Fe-2S] clusters, and directly into enzymes^25,26^. 10 DPT cells are reminiscent of PCBP1 deficient models that have poor ferritin loading, increased labile iron pool and lipid peroxidation^23,34^. Loss of PCBP1 in surviving cells likely explains the counterintuitive vulnerability to ferroptosis when the NRF2, GSH, Ferritin axis is enriched. High levels of GSH in cells 10 DPT is needed to both bind excess labile iron and manage redox requirements, including regeneration of the GPX4 active site, allowing cells 10 DPT to avoid spontaneous ferroptotic cell death. However, without attenuating the source of lipid peroxidation, i.e., high labile iron, surviving cells are reliant on lipid peroxidation defenses and susceptible to GPX4 inhibition. Deficient iron trafficking from PCBP1 loss may also be impacting the function of other iron containing enzymes in cells 10 DPT, particularly in the mitochondria and nucleus, and contributing to the convergent transcriptomic signature. In addition to chaperoning iron, PCBP1 has separate roles in RNA processing and has been implicated for gene regulation in epithelial-mesenchymal transition and innate immune signaling, as well as binding oxidized RNA^49–51^. While it is tempting to speculate that loss of PCBP1 is related to RNA metabolism imbalances or activation of one pathway enabling chemotherapy survival, while consequently contributing to labile iron accumulation, more research is needed to tease out PCBP1’s distinct and independently vital roles^33^.

Dysregulation of cellular iron homeostasis is causal or characteristic to many diseases, and ferroptosis is an increasingly appreciated contributor to many pathologies^41,52–54^. Labile iron accumulation in cells 10 DPT was visibly punctate compared to untreated cell lines, with much of the punctate staining colocalizing with active lysosomes. Lysosomal accumulation of iron is characteristic to chronic stress conditions, particularly neurodegenerative diseases, but whether iron sequestration in lysosomes is protective or not is unclear. On the one hand, containing iron in lysosomes prevents diffusion and labile irons access to other cellular compartments. However, lysosomal iron is not detoxified as it is in ferritin, and thus causes lipid peroxidation, increased lysosomal permeability, and oxidation of proteins that accumulate in insoluble aggregates^20,41,54^. In regard to ferroptosis, lysosomal iron is known to be essential for ferroptosis execution due to deferoxamine’s ability to prevent ferroptosis in most contexts^3^. Indeed, deferoxamine prevented ferroptotic cell death by GPX4 inhibition in all samples in this study (Supplementary Fig. S11). Additionally, a recent study showed that lysosomal iron can trigger ferroptosis by initiating lipid peroxidation in lysosomes that then propagate to ER and other intracellular membranes^55^. Intracellular localization of labile iron remains a critical underexplored area of ferroptosis research, and lysosomal ferrous iron pools constitute a novel target for increasing sensitivity to ferroptosis in resistant cancer settings^55,56^.

Ferroptosis is an emerging vulnerability in resistant cancer settings. In our study, cells surviving chemotherapy are also vulnerable to ferroptosis, but this vulnerability is driven by labile iron accumulation, partially in lysosomal compartments, and occurs in spite of active NRF2 signaling. An inability of ferritin to relieve labile iron burden is also counterintuitive. This study reinforces the highly flexible mechanisms involved in ferroptosis and the importance of timing when considering therapeutic strategies^9^. Iron homeostasis dynamics in cancer, intracellular localization of reactive iron, and the impact of PCBP1 on intracellular iron handling, are important areas of further research to enable translation of ferroptosis therapeutic strategies.

## METHODS

### Cell culture and generation of surviving cells

PC3 cells were cultured in RPMI (Gibco #11875119) with 10% fetal bovine serum (FBS, Avantor #97068-085) and 1% penicillin–streptomycin. MDA-MB-231 cells were cultured in DMEM (Gibco #11995073) with 10% FBS and 1% penicillin–streptomycin. Surviving cells were generated by treating wish IC50 cisplatin or docetaxel for 72 hours, then chemotherapy was removed and cells maintained in fresh media with routine passaging. PC3 chemotherapy treatment was 6uM or 5nM docetaxel. MDA-MB-231 treatment was 12uM cisplatin or 5nM docetaxel.

### Single cell RNA sequencing and data processing

Cells were collected on timepoint day using tubes pre-coated with bovine serum albumin (BSA). Cell fixation was performed on timepoint day using Parse Biosciences Fixation kit (Parse Biosciences) according to manufacturer protocol, except for no cell filtering steps were performed. No cell filtering steps were performed during barcoding and library prep to prevent loss of the large cells surviving chemotherapy. Barcoding and library prep for MDA-MB-231 cells untreated and surviving cisplatin or docetaxel was done using the Evercode Whole Transcriptome Mega kit (V2, Parse Biosciences). Sequencing was performed on an Illumina NovaSeq 6000 and alignment done using v0.7.2p alignment pipeline (Parse Biosciences). Barcoding and library prep for PC3 cells untreated and surviving cisplatin was done using the Evercode Whole Transcriptome kit (V2, Parse Biosciences). Sequencing was performed on an Illumina NovaSeq 6000 and alignment done using v0.7.2p alignment pipeline (Parse Biosciences). Barcoding and library prep for PC3 cells untreated and surviving docetaxel was done using the Evercode Whole Transcriptome kit (V2, Parse Biosciences). Sequencing was performed on an Illumina NovaSeq X and alignment done using v1.0.6p alignment pipeline (Parse Biosciences).

RNA barcodes were filtered on Total UMI Count (> 500 UMIs), feature count (> 350 features), and percentage of mitochondrial genes (< 25%). Seurat v4.3.0^57^ was used for handling of normalization, identification of variable genes, scaling, principal component analysis, UMAP dimensional reduction, and SNN generation followed by Leiden clustering. Batch effect correction was performed using Harmony^58^. Clusters were identified using a combination of marker genes and differential expression comparing each cluster to all other cells in the data set. Differential expression analyses were performed using Mann-Whitney U test and fgsea v1.24.0 was used to run gene set enrichment analysis (GSEA). Features were ranked by -log(pvalue) * sign(foldchange). Percentages of cells by cluster and sample are the percentage of cells in each cluster compared to the total number of cells per sample. Statistical comparisons shown are from t-test comparing the percentages.

For PC3 cells, cisplatin treated and docetaxel treated samples were collected and sequenced separately, both with an untreated sample. To avoid batch effect contribution, all GSEA was performed only compared to the respective control (untreated) sample and Ucell depictions are separated by chemotherapy and include only the respective control (untreated) sample on each plot. Genesets used for initial GSEA and UCell were from the Molecular Signatures Database^59^ Hallmark and Kegg collections (Fig. 1, Fig. 2A, B). Iron handling genesets were from the Molecular Signatures Database Gene Ontology collection (Fig. 2C). Autophagy related genesets are from literature. ‘TFEB Signaling’ is from^60^ Table S2 TFEB-KO vs TFEB-WT gene expression list, genes less than −1.5 LogFC (193 genes). ‘Autophagy’ (Table S1) is from^61^ Table S1 ‘autophagy core’ excluding ‘its mutation leads to autophagy defects’ and ‘autophagy dependent cell death’ and ‘negative regulators’. ‘Autophagy Induction’, and ‘Lysosome Biogenesis’ are also from^61^ Figure 2.

### Drug treatments

Drugs used in this study are Bafilomycin (SelleckChem #S1413, 1uM); deferoxamine mesylate (SelleckChem #S5742, 50uM, DFO); Ferrostatin-1 (SelleckChem #S7243, 1uM, Fer-1); ML210 (SelleckChem #S0788, 1uM).

### Labile iron live cell imaging

Cells were lifted using TypLE Express (Gibco #12604039) and re-plated on to 35 mm IbiTreat µ-Dishes (Ibidi USA Inc., #81156). Untreated cell lines were seeded the day prior to imaging. For 1 DPT, cells were collected and re-plated at the end of chemotherapy treatment. For 5 DPT cells, cells were collected and re-plated at 3 DPT. For 10 DPT cells, cells were collected and re-plated at 7 DPT. On timepoint day, dishes were washed once with HBSS (Gibco #14025-092) and then incubated with FerroOrange (Millipore Sigma #SCT210, 1uM); ER-Tracker Green (BODIPY™ FL Glibenclamide) (Invitrogen #E34251, 1uM); and Hoechst 33342 (Invitrogen #I35101B, 1uM) in HBSS for 30 minutes at 37°C. Dishes were then washed twice with HBSS and left in HBSS for imaging. Imaging was done one dish at a time, with no more than 30 minutes elapsing before completion of imaging. Images were acquired using a Zeiss Observer Z1 microscope/ZEN pro 2.0 software (Carl Zeiss Microscopy) with a Axiocam MRm camera (Zeiss) and X-Cite Xylis LED Fluorescence Light Source (Excelitas Technologies). Images were background subtracted via Rolling Ball background subtraction, manual ROIs drawn per cell using phase and Hoechst imaging, and then integrated density measured per cell using Fiji^62^. For lysosomal colocalization, FerroOrange staining was the same as described, plus LysoTracker Deep Red (Invitrogen #L12492, 50nM) was spiked into the final 15 minutes of staining time period. Image acquisiton was the same as described using higher magnification objective. Colocalization analysis was performed using Fiji.

### Labile iron assay from lysates

On timepoint day, cells were lifted using TypLE Express, washed once in HBSS, counted with hemacytometer and then snap frozen. Assay was run no more than 7 days after snap freezing. For assay, cell pellets were lysed using Mammalian Cell Lysis Buffer 5X (Abcam #ab179835) with Halt Protease and Phosphatase Inhibitor Cocktail (Thermo Scientific #78440) added. Iron Assay Kit (colorimetric) (Abcam #ab83366) was then run according to manufacturer protocol.

### Western Blot

On timepoint day, cells were lifted using Cell Dissociation Solution Non-enzymatic 1x (Millipore Sigma #C5914), washed once in PBS and then snap frozen. Cell pellets were lysed no more than 7 days after snap freezing. Cell pellets were lysed using RIPA buffer with Halt Protease and Phosphatase Inhibitor Cocktail (Thermo Scientific #78440). Protein concentration was determined by bca assay. Western blot samples were prepared using Laemmli Sample Buffer (Bio-Rad) supplemented with β-mercaptoethanol (Bio-Rad). The same total protein amounts were loaded into each lane of 4–20% Mini-PROTEAN TGX Stain-Free Protein Gels (Bio-Rad). Completed gels were transferred onto 0.2 um nitrocellulose membrane. Membranes were blocked using Casein Blocking Buffer (Millipore Sigma #B6429) and then incubated overnight in primary antibodies diluted in blocking buffer. Membranes were washed using Tris-buffered saline with 0.1% Tween 20 after incubation in both primary and secondary antibodies. Membranes were imaged using Li-COR Odyssey system. Blots were quantified in Fiji with each target protein densitometry normalized to respective actin loading control densitometry. Antibodies used in this study were B-actin (Sigma-Aldrich #A5441, RRID:AB_476744, 1:5000); Ferritin (Abcam #ab75973, RRID:AB_1310222, 1:800); Transferrin Receptor (Abcam #ab269513, RRID:AB_3351672, 1:3000); NRF2 (Abcam #ab62352, RRID:AB_944418, 1:800); PCBP1 (Abcam #ab168377, RRID: AB_3665910, 1:1000); GPX4 (Abcam #ab125066, RRID:AB_10973901, 1:1000); LC3B (Cell Signaling Technology #3868, RRID:AB_2137707, 1:1000); p62 (Cell Signaling Technology #39749, RRID:AB_2799160, 1:1000); IRDye 680RD Goat anti-Mouse IgG Secondary Antibody (LICOR Biosciences #926-68070, RRID:AB_10956588, 1:20000); IRDye 800cw Goat Anti-Rabbit IgG Secondary Antibody (LI-COR Biosciences #926-32211, RRID:AB_621843, 1:12000).

### GSH / GSSG assay

Cells were plated into 96 well plates prior to assay being run (as described in ‘Labile iron live cell imaging’ section). GSH / GSSG ratio was determined using GSH/GSSG-Glo Assay kit (Promega #V6611) according to manufacturer protocol. Both untreated cancer cell lines were included in every assay plate as a control. Any wells that were > 80% confluent were not assayed. All reagents were made prior to washing and on plate lysis as described in protocol to ensure rapid assay completion and as accurate a determination of GSH/GSSG ratio as possible.

### C11 Bodipy analysis

Cells were seeded into 24 well plates (as described in ‘Labile iron live cell imaging’ section). C11 BODIPY 581/591 (Invitrogen# D3861, 1uM) dissolved in HBSS was added to each sample. Cells were incubated for 30 minutes at 37°C, washed once in HBSS, then imaged in HBSS. Images were acquired using an EVOS M7000 imaging system. Images were quantified in Fiji by manually drawing ROIs per cell using phase and Hoechst images. Red and green fluorescence values were background-corrected by quantifying the red or green fluorescence in cell-free areas from at least four cell free ROIs and subtracting from per cell values to determine the final fluorescence values. C11 Bodipy ratio is green fluorescence (oxidized) / red + green fluorescence total).

### Ferroptosis cell death assay

Cells were seeded into 6 well plates (as described in ‘Labile iron live cell imaging’ section). Cells were treated with DMSO, ML210, ML210 and Fer-1, or ML210 and DFO in DMEM supplemented with Incucyte Cytotox Green Dye (Sartorius #4633). Live cell timelapse microscopy was performed using an Incucyte SX5 (Sartorius) with imaging every hour for 10 hours. Movies were assembled in Fiji and manually counted to determine cell loss (detachment from apoptosis) and cell death by ferroptosis (adherent, but positive for Cytotox Green, at 10 hours). Every plate included one well for each condition, and 100 ∼ 150 cells were counted per well across at least two fields of view. Ferroptosis confirmed by Fer-1 and DFO fully preventing ferroptotic cell death.

### Release of immunomodulatory markers during ferroptosis

Immunomodulatory molecule detection experiments were performed at Cellomatics Biosciences, 10 Colwick Quays Business Park, Private Road No.2, Colwick, Nottingham NG4 2JY, United Kingdom.

Cells were seeded into 96 well plates as described. On assay day, cells were treated with DMSO or ML210 for 6 hours prior to co-culture with PBMCs. Extracellular ATP release using the RealTime-Glo Extracellular ATP Assay (Promega #GA5010) according to the manufacturer’s protocol. HMGB1, M-CSF, and IL-2 were detected via ELISA (ThermoFisher #EEL047) and Luminex Multiplex Assay (R&D #FCSTM18B-10).

PBMCs were isolated as follows. Three healthy blood donors were recruited, and fresh whole blood was collected on the day of co-culture for each timepoint into BD Vacutainer tubes coated with EDTA (BD #367525). PBS was added to the whole blood collected in EDTA coated vacutainer tubes at equal volumes, and tubes were mixed gently. Approximately 15 mL of Lymphoprep (StemCell #7801) was added into SepMate tubes (StemCell #85450), which was kept vertical in a tube rack. The diluted blood sample was then gently pipetted down the sides of the tubes. The tubes were centrifuged at 1200 g for 12 minutes at room temperature. The PBMCs above the plastic insert of the SepMate tube were transferred into a 50 mL tube. PBMCs were washed with 10 mL PBS. The tubes were centrifuged at 1200 g for 12 minutes at room temperature. The PBMC pellets were resuspended in 1X RBC lysis buffer and incubated at room temperature for 10 minutes before being centrifuged at 600 g for 7 minutes. The PBMC pellets from each donor were then resuspended in pre-warmed complete media (the same media used for cells) and counted based on trypan blue dye exclusion. Equal PBMC numbers were pooled from each donor for the co-cultures.

### Statistical Analysis

All statistical analyses were performed in GraphPad Prism version 9.1 (GraphPad Software, LLC). An α = 0.05 (confidence level 95%) was the criterion considered to determine statistical significance for all tests. No significance (ns), *, **, ***, and **** represent p values of ≥0.05, <0.05, <0.01, <0.001, and <0.0001, respectively.

