## Supplementary Figures for "Dysregulation of cellular iron predisposes chemotherapy resistant cancer cells to ferroptosis"

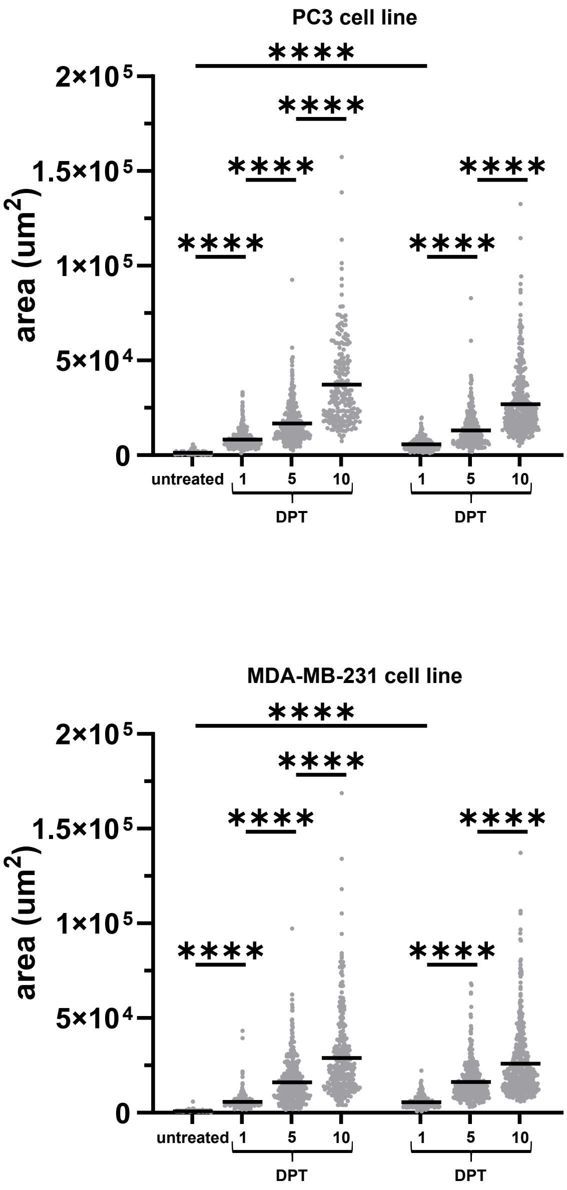


**Supplementary Figure S1 Cells surviving chemotherapy increase in size over time.**
Area of PC3 and MDA-MB-231 cell lines before and after cisplatin or docetaxel as determined by live cell phase contrast imaging.

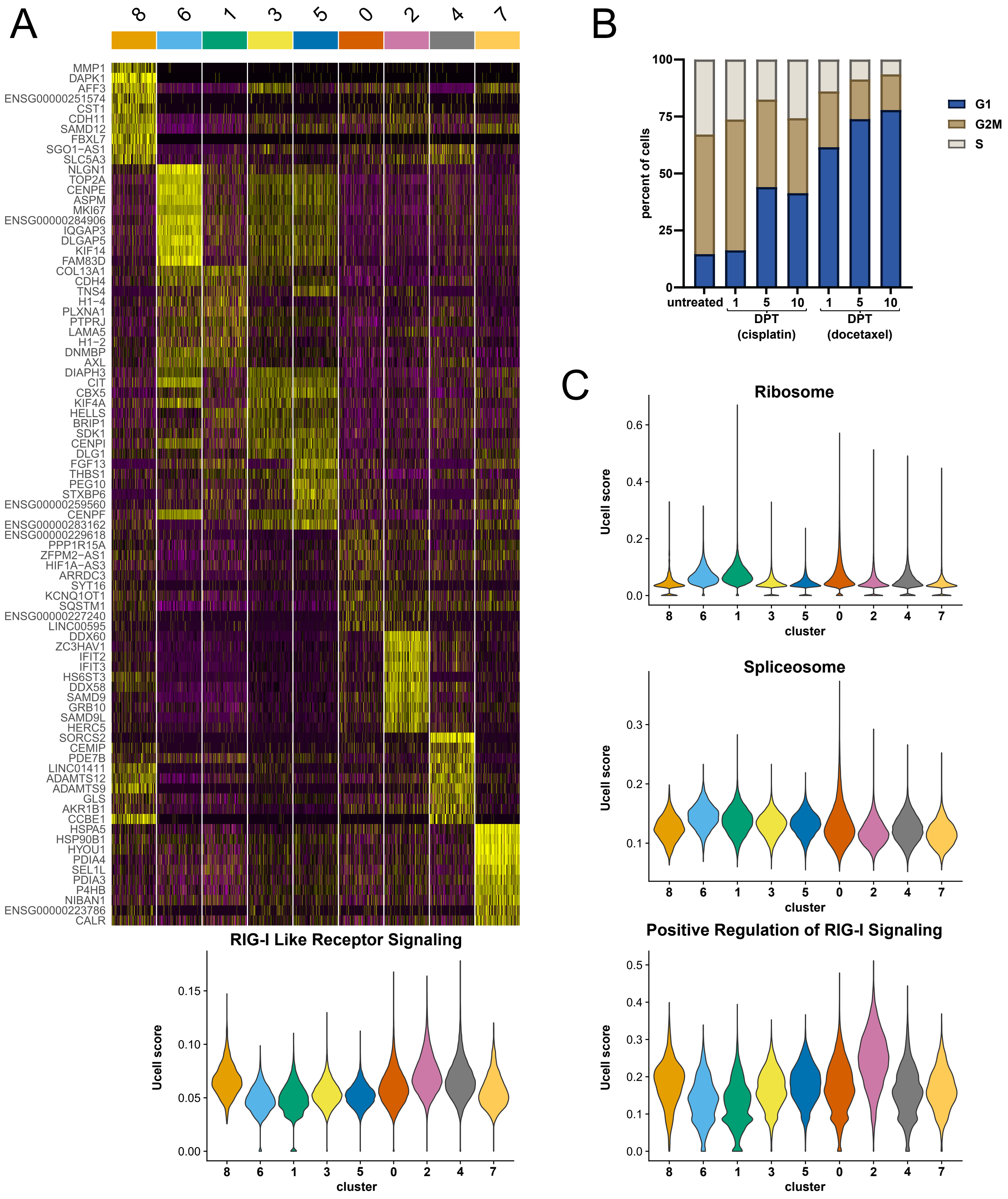


**Supplementary Figure S2 Heterogeneity of MDA-MB-231 cells transcriptional response to chemotherapy. A.** Cluster defining genes for UMAP projection of MDA-MB-231 cells treated with cisplatin or docetaxel (Fig. 1A). **B.** Cell cycle scoring for surviving cells at different DPT. **C.** Ucell score of selected genesets across each cluster in Fig 1A.

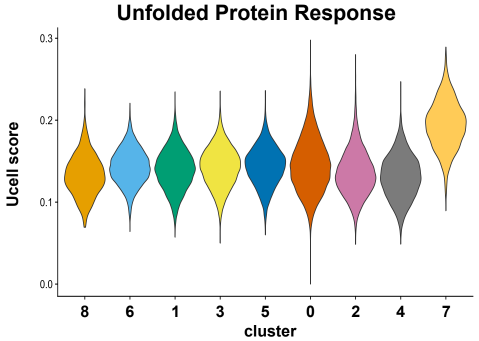

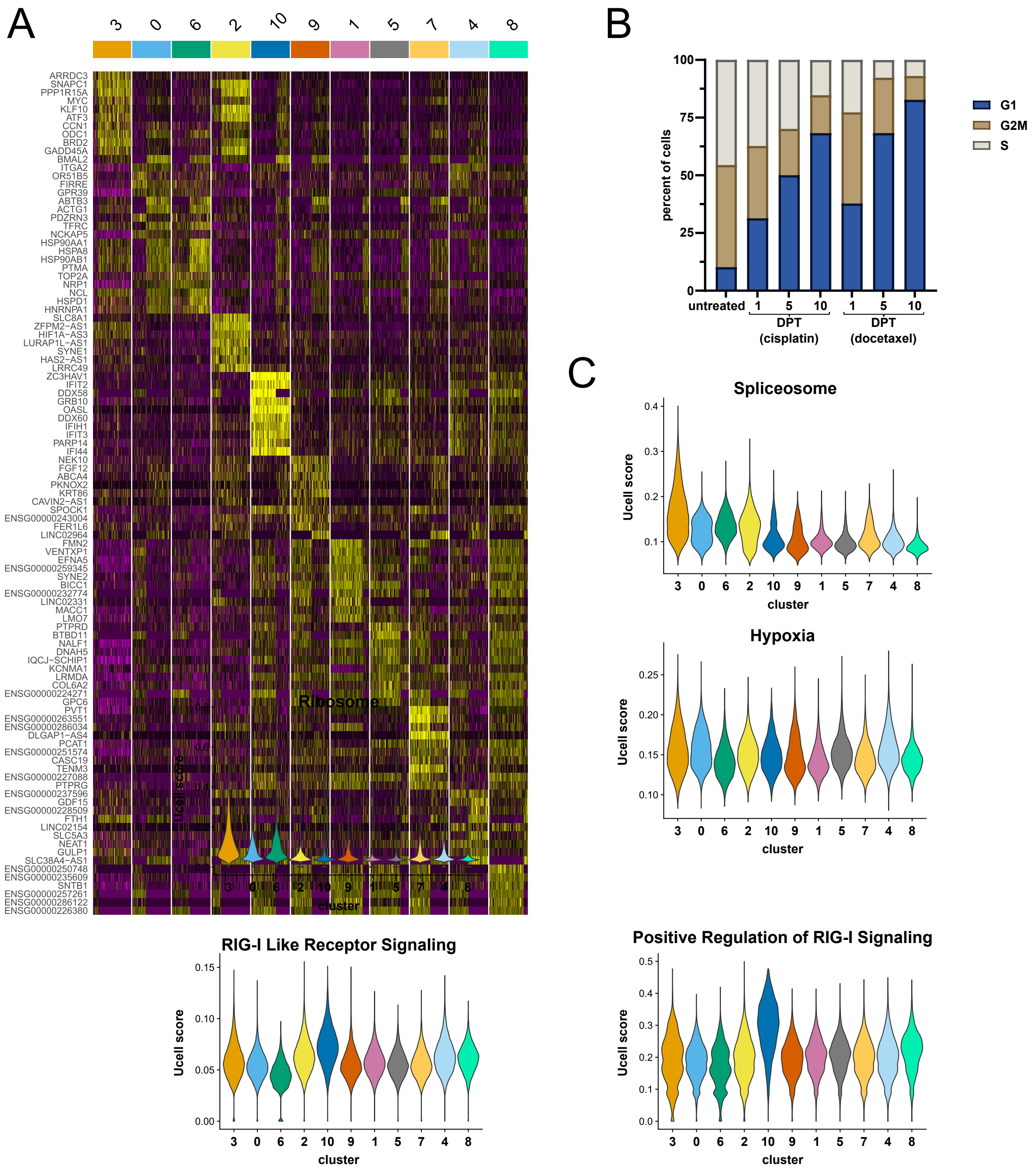


**Supplementary Figure S3 Heterogeneity of PC3 cells transcriptional response to chemotherapy. A.** Cluster defining genes for UMAP projection of PC3 cells treated with cisplatin or docetaxel (Fig. 1D). **B.** Cell cycle scoring of surviving cells across DPT. **C.** Ucell score of selected genesets across each cluster in Fig 1D.

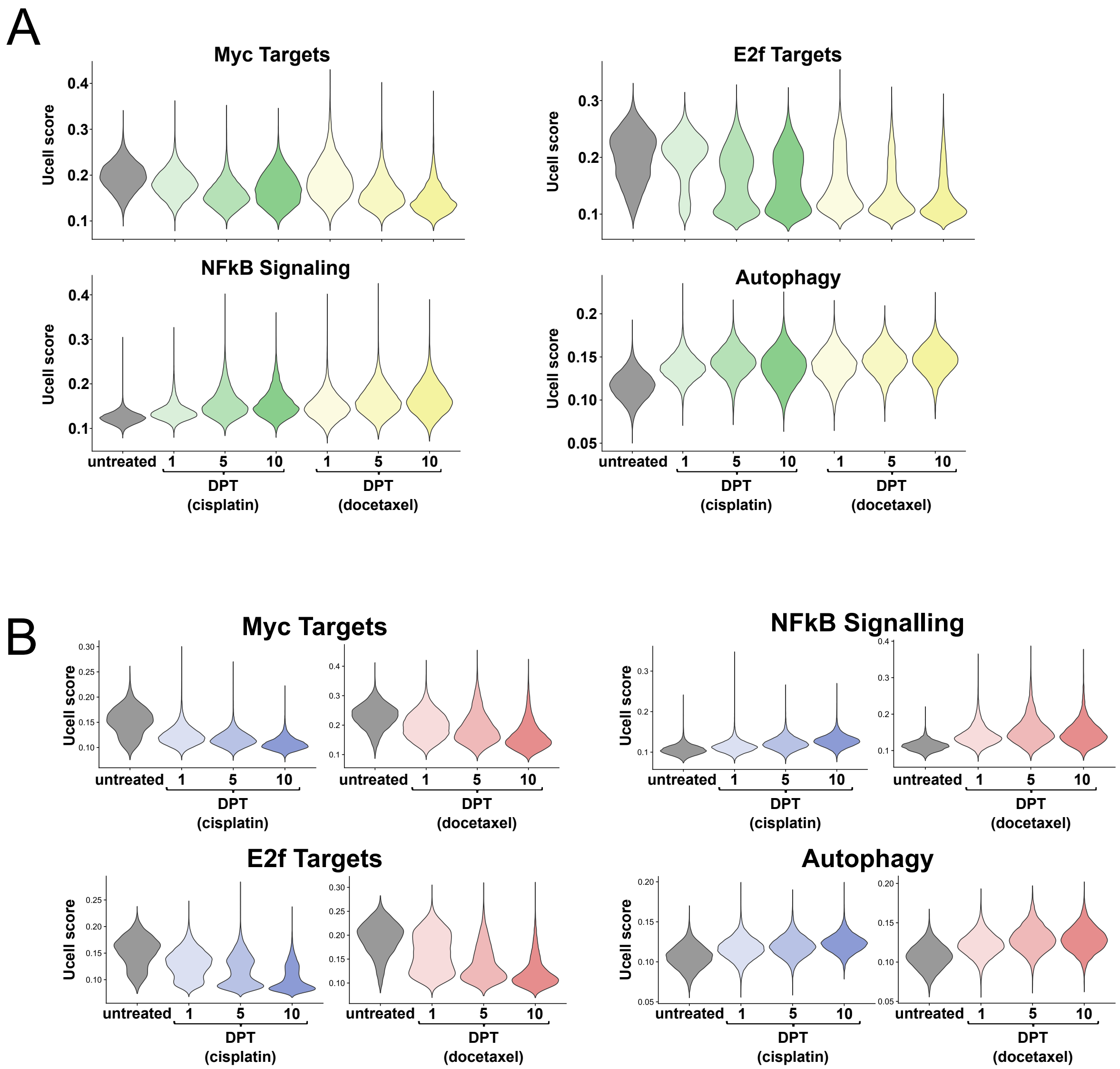


**Supplementary Figure S4. Transcriptomic signatures of cells surviving chemotherapy converge over time. A.** Ucell scoring for select genesets in MDA-MB-231 cells surviving chemotherapy. Violin plot depiction of all cells within each sample. **B.** Ucell scoring for select genesets in PC3 cells surviving chemotherapy. Violin plot depiction of all cells within each sample.

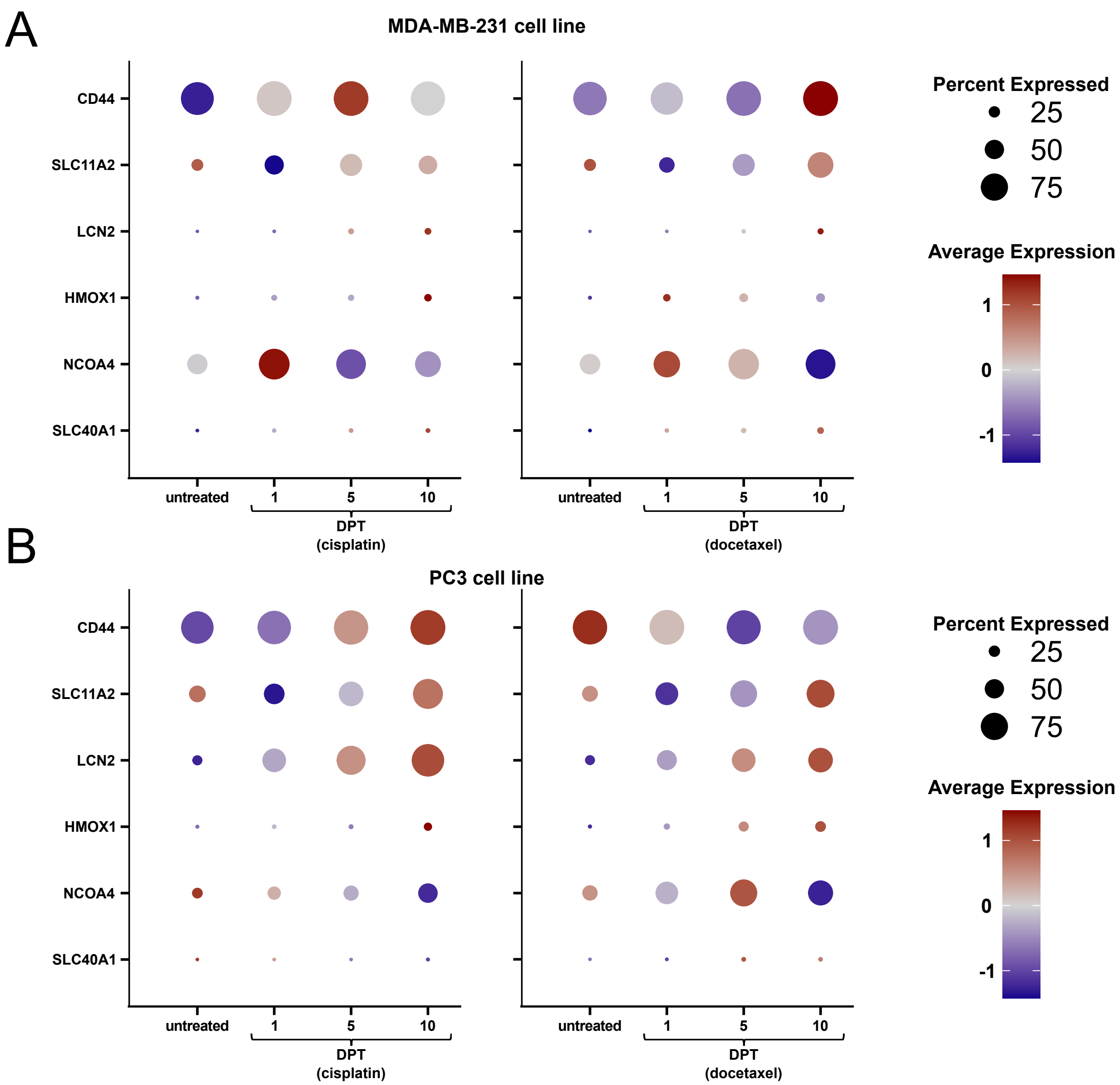
A
MDA-MB-231 cell line
CD44
SLC11A2
LCN2
HMOX1
NCOA4
SLC40A1
untreated
1
5
10
DPT
(cisplatin)
DPT
(docetaxel)
Percent Expressed
25
50
75
Average Expression
1
0
-1
B
PC3 cell line
CD44
SLC11A2
LCN2
HMOX1
NCOA4
SLC40A1
untreated
1
5
10
DPT
(cisplatin)
DPT
(docetaxel)
Percent Expressed
25
50
75
Average Expression
1
0
-1

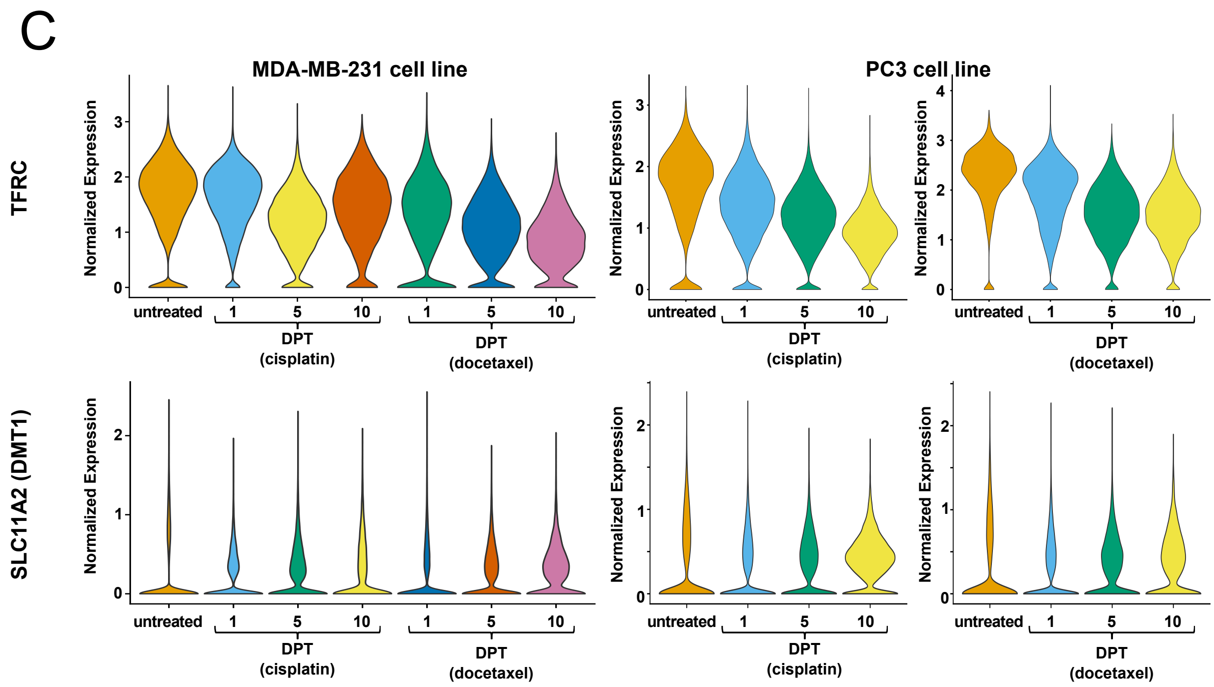


**Supplementary Figure S5 Gene expression of iron homeostasis genes in cells surviving chemotherapy.** Iron homeostasis genes in MDA-MB-231 (**A**) or PC3 (**B**) cells surviving cisplatin or docetaxel. Each gene represented by z-scored distribution of average normalized expression across samples (color) and percentage of cells with detected transcript (size). **C.** Normalized expression of Transferrin Receptor (TFRC) and DMT1 (SLC11A2) in MDA-MB-231 and PC3 cells surviving cisplatin or docetaxel.

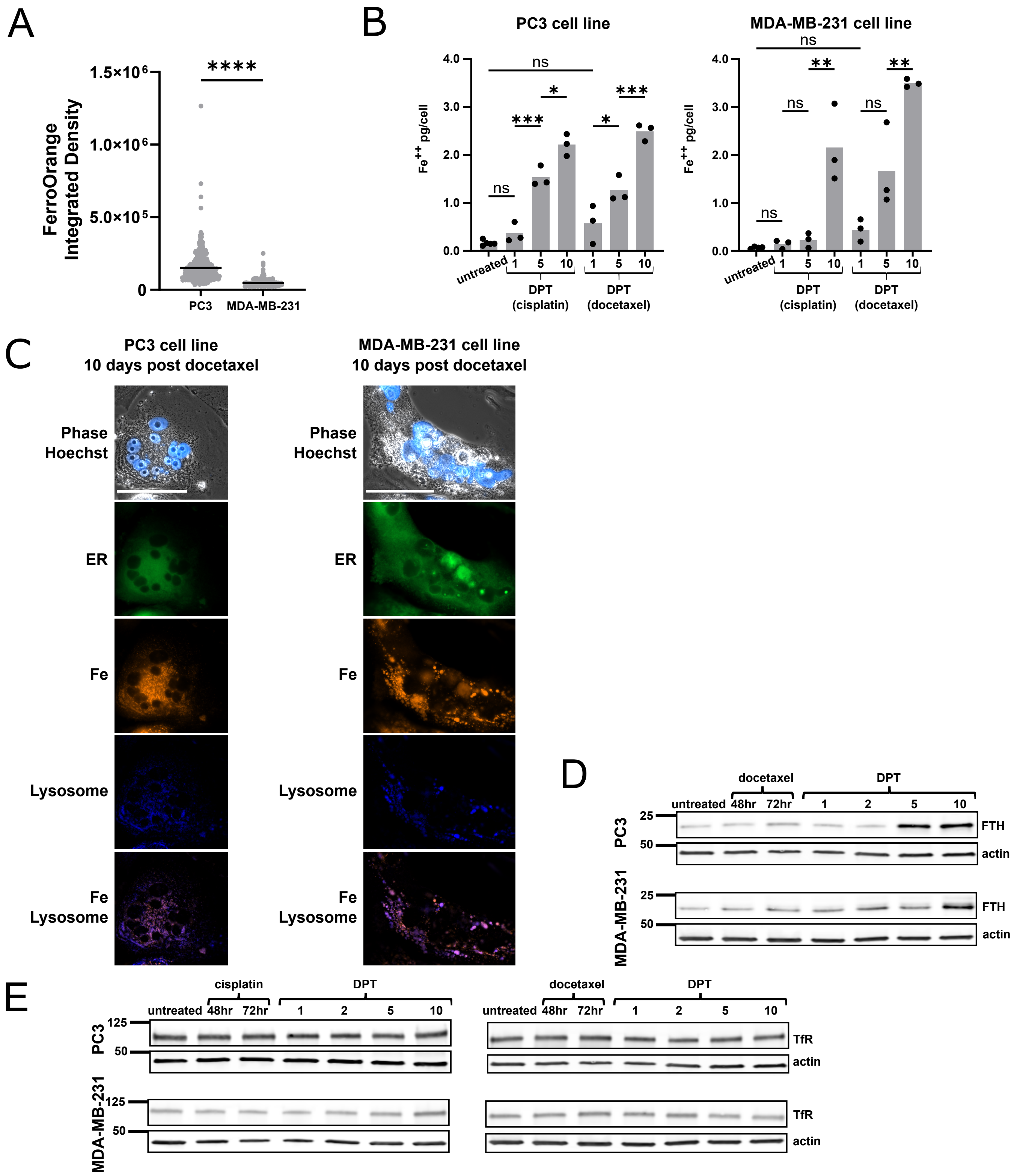


**Supplementary Figure S6. Labile iron is dramatically elevated by 10 days recovery from chemotherapy. A.** Quantification of FerroOrange integrated density in untreated cell lines from Fig. 3A. Data displays individual cells (dot) and mean (bar) from three biological replicates **B.** Labile iron assay from cell lysates of PC3 and MDA-MB-231 cells before and after chemotherapy. **C.** Live cell images with FerroOrange (Fe), ERTracker Green (ER), and LysoTracker (Lysosome) of PC3 and MDA-MB-231 cells 10 days after cisplatin. Quantification in Figure 3D. **D.** Ferritin expression in PC3 and MDA-MB-231 cells before and after docetaxel. **E.** Transferrin Receptor expression in PC3 and MDA-MB-231 cells before and after cisplatin or docetaxel.

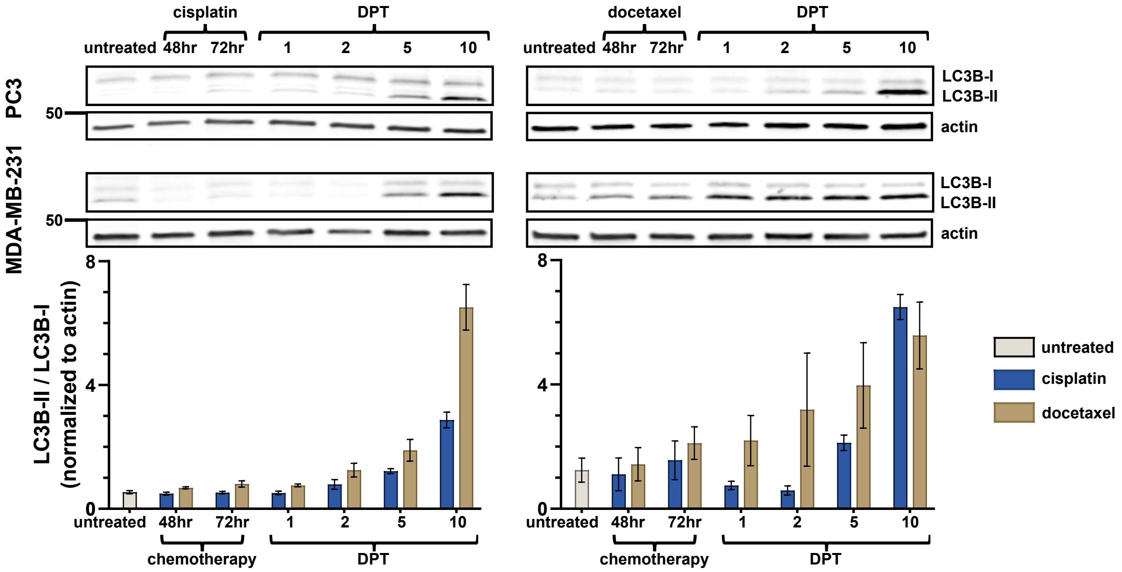


**Supplementary Figure S7. Transcriptional and protein enrichment of autophagy in cells surviving chemotherapy. A.** Western blot for LC3B in PC3 or MDA-MB-231 cells surviving cisplatin of docetaxel. Quantification is LC3B-II / LC3B-I ratio.

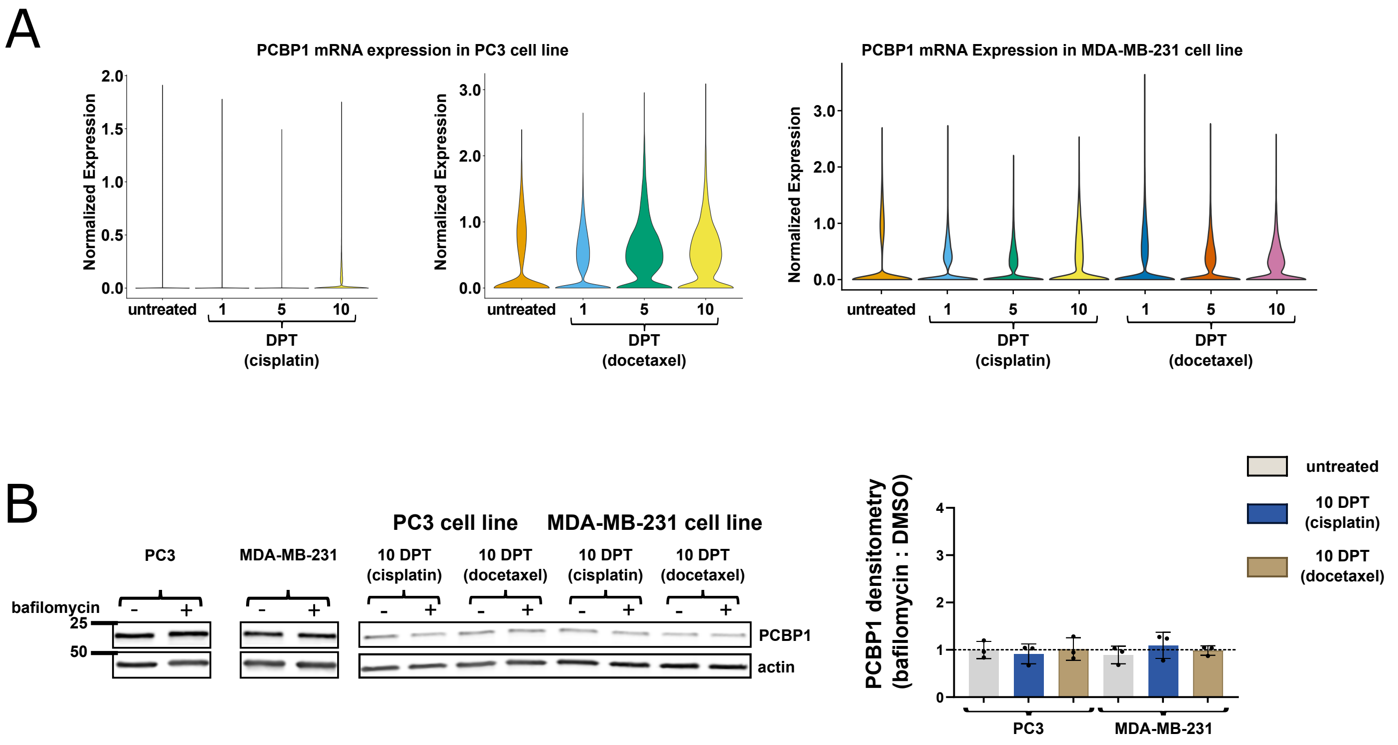


**Supplementary Figure S8 Loss of PCBP1 in cells surviving chemotherapy is not due to transcriptional regulation or autophagic degradation.**

**A.** PCBP1 RNA expression in PC3 or MDA-MB-231 cells surviving cisplatin or docetaxel.

**B.** Western blot for PCBP1 expression following 1 uM bafilomycin or DMSO treatment for 6 hrs. Quantification shows bafilomycin : DMSO treated densitometry ratio. Actin bands same as in Fig. 4E.

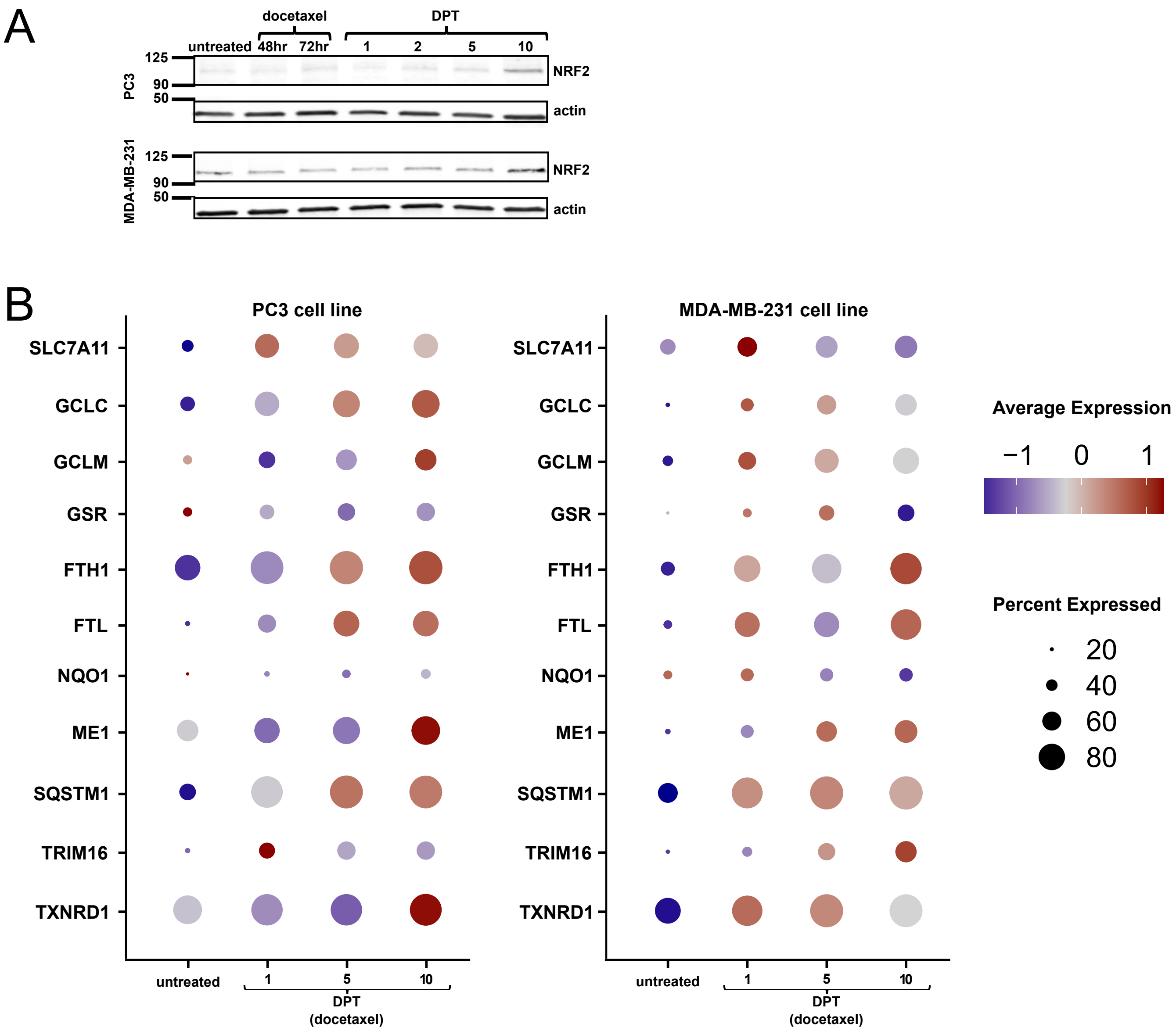
A
docetaxel
DPT
untreated
48hr
72hr
1
2
5
10
125
90
50
PC3
NRF2
actin
MDA-MB-231
125
90
50
NRF2
actin
B
PC3 cell line
MDA-MB-231 cell line
SLC7A11
GCLC
GCLM
GSR
FTH1
FTL
NQO1
ME1
SQSTM1
TRIM16
TXNRD1
untreated
1
5
10
DPT
(docetaxel)
Average Expression
−1
0
1
Percent Expressed
20
40
60
80

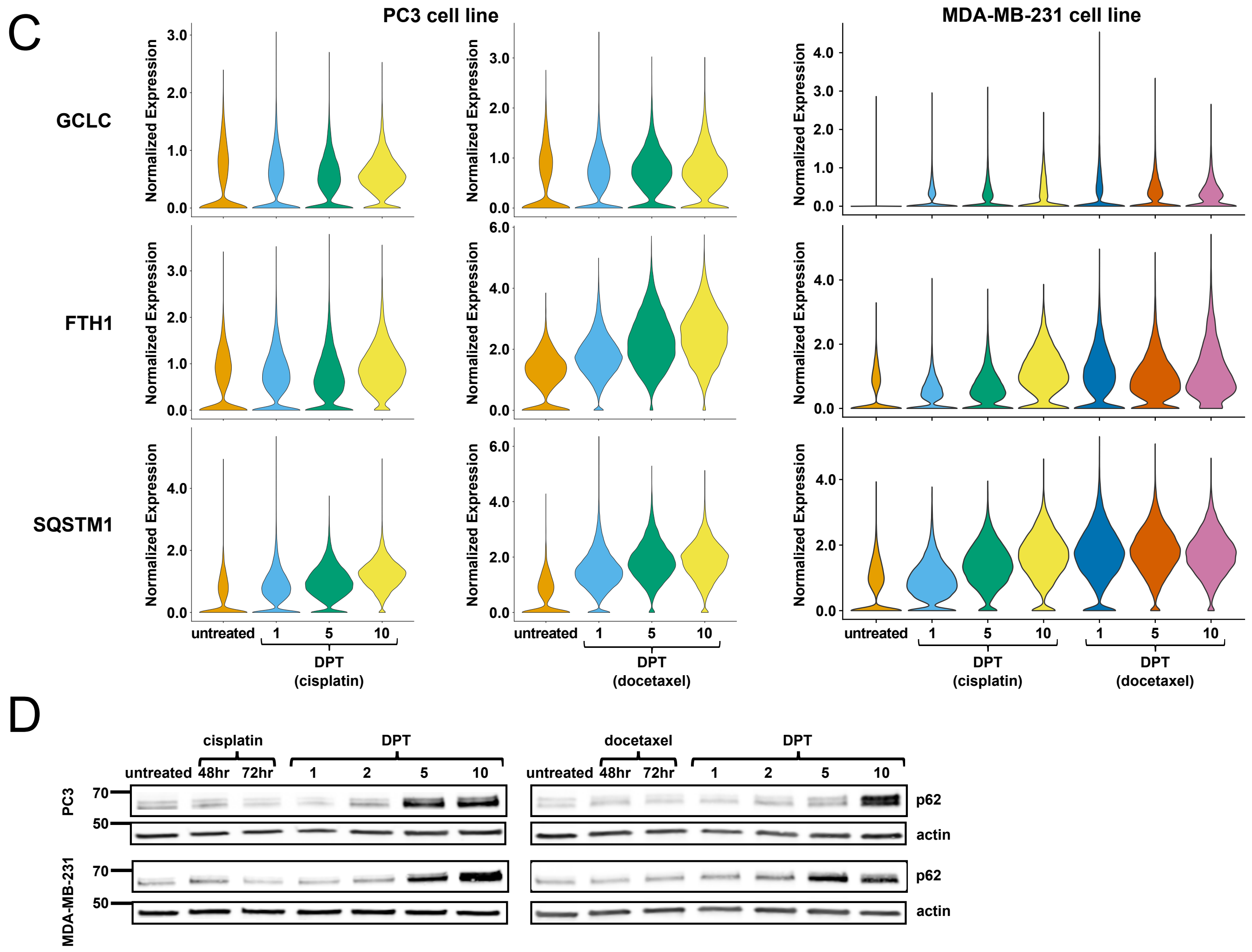


**Supplementary Figure S9. Surviving cells 10 days post chemotherapy have active NRF2 signature. A.** Western blot for NRF2 in PC3 and MDA-MB-231 cells before and after docetaxel. **B.** NRF2 target gene expression in PC3 or MDA-MB-231 cells surviving docetaxel. Each gene represented by z-score distribution across samples (color) and percentage of cells with detected transcript (size). **C.** Normalized expression of NRF2 target genes displayed as violin plot of all cells per sample. **D.** Western blot for p62 expression in PC3 and MDA-MB-231 cells following cisplatin or docetaxel treatment.

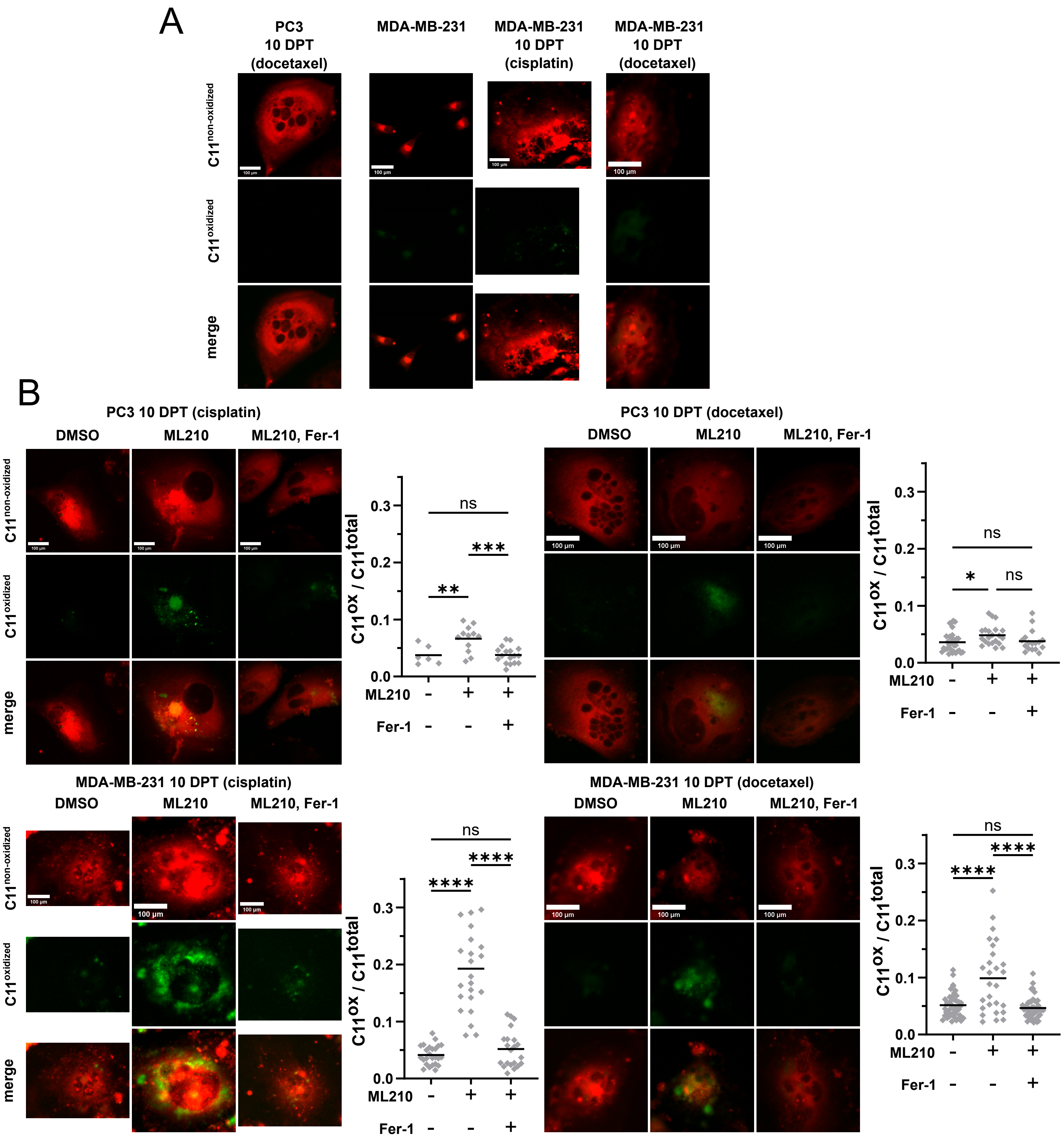


**Supplementary Figure S10 GPX4 inhibition leads to a build up of oxidized lipids.** **A.** C11 Bodipy imaging in PC3 cells 10 days post docetaxel, and MDA-MB-231 cells untreated and 10 days post cisplatin or docetaxel. Quantification in 6A. **B.** C11 Bodipy imaging and quantification for all 10 DPT surviving cell contexts treated with DMSO, ML210, or ML210 and Ferrostatin-1 (Fer-1) treatment for 6 hours. Quantification is ratio of oxidized (green) to total (green + red) C11 Bodiy fluorescence.

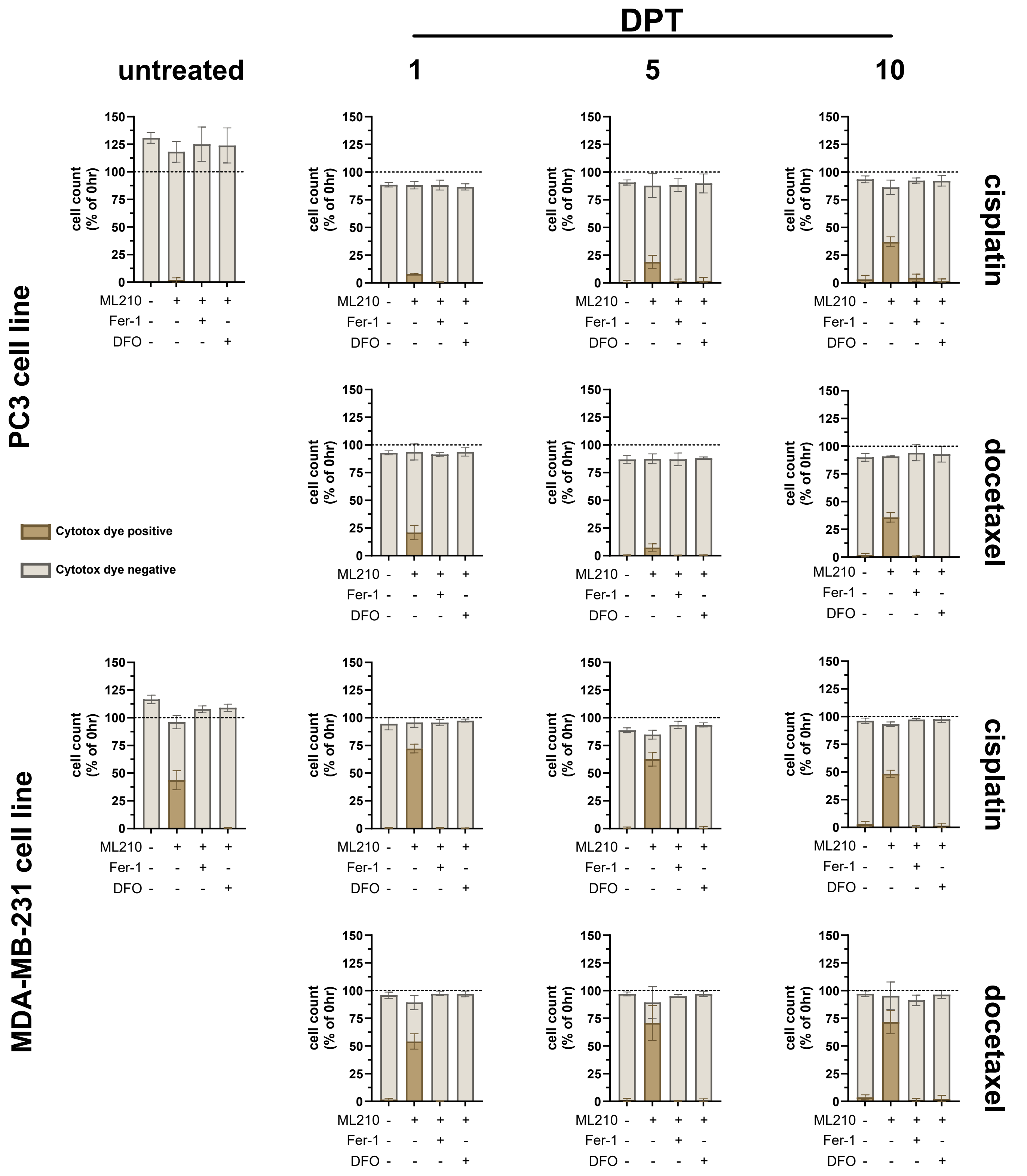


**Supplementary Figure S11 Ferrostatin-1 and Deferoxamine prevent ferroptotic cell death from GPX4 inhibition.** All untreated cell lines and timepoints of surviving cells following chemotherapy treated with, from left to right, DMSO (0.1%); ML210 (1uM); ML210 and Fer-1 (1uM); ML210 and DFO (50uM). Percent of total cell count is change in cell number over 10 hours from cell detachment (apoptosis) or cell proliferation. Cytotox Green dye positive cells are still adherent at end of timelapse (10 hours). PC3 cells 10 DPT (cisplatin) are repeated from Figure 6D.

Fer-1 = Ferrostatin-1  DFO = Deferoxamine

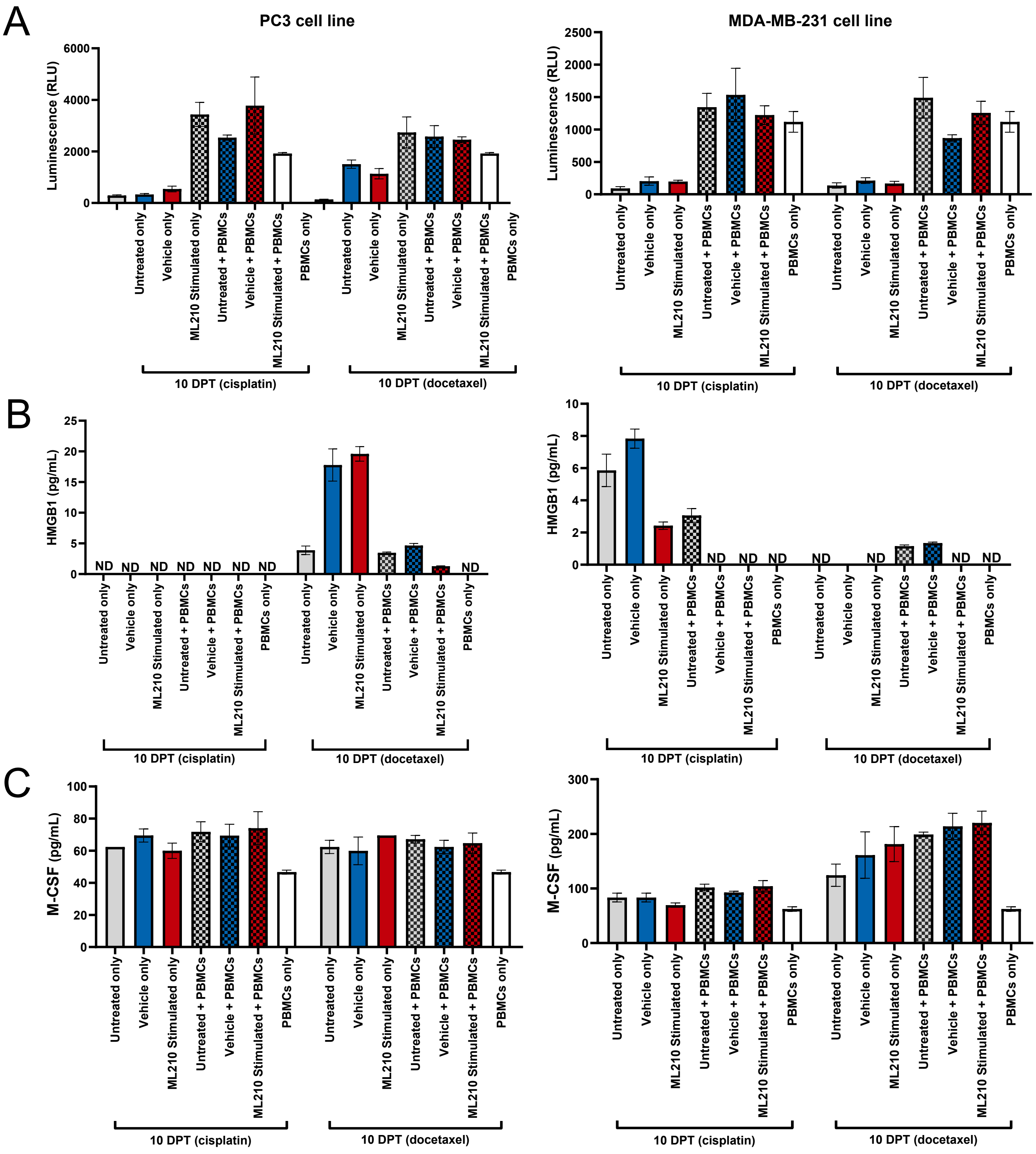

A
PC3 cell line
MDA-MB-231 cell line
Luminescence (RLU)
B
HMGB1 (pg/mL)
ND
C
M-CSF (pg/mL)
Untreated only
Vehicle only
ML210 Stimulated only
Untreated + PBMCs
Vehicle + PBMCs
ML210 Stimulated + PBMCs
PBMCs only
10 DPT (cisplatin)
10 DPT (docetaxel)

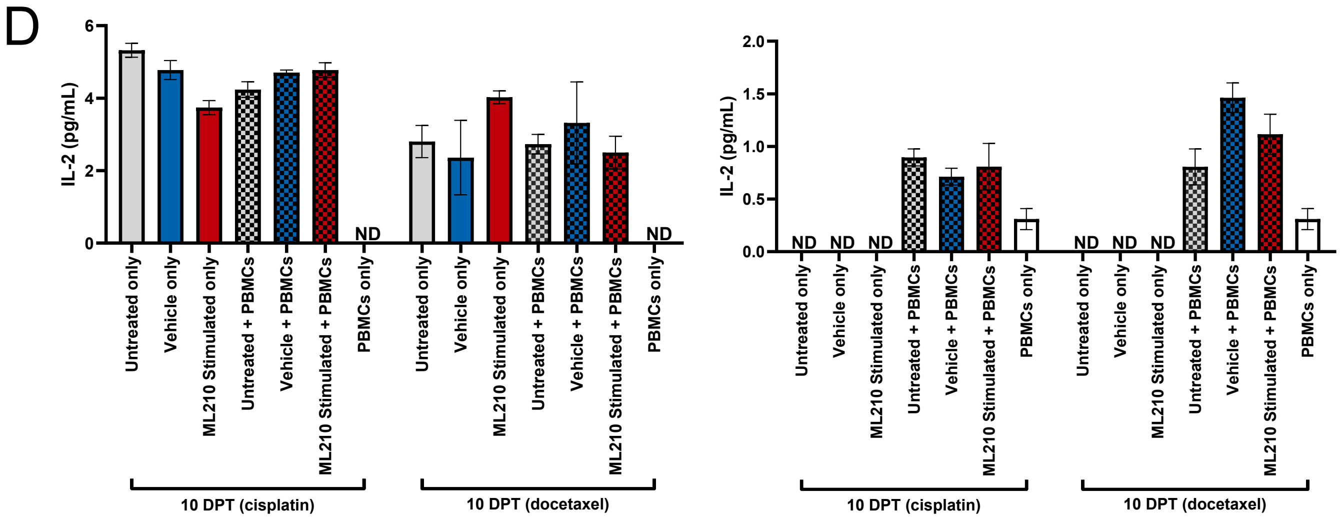


**Supplementary Figure S12 Ferroptotic cell death does not release immunogenic molcules. A.** PC3 or MDA-MB-231 surviving cells 10 days post chemotherapy and treated with ML210 were assessed for ATP (**A**), HMGB1 (**B**), M-CSF (**C**), and IL-2 (**D**) in presence or absence of human PBMCs.

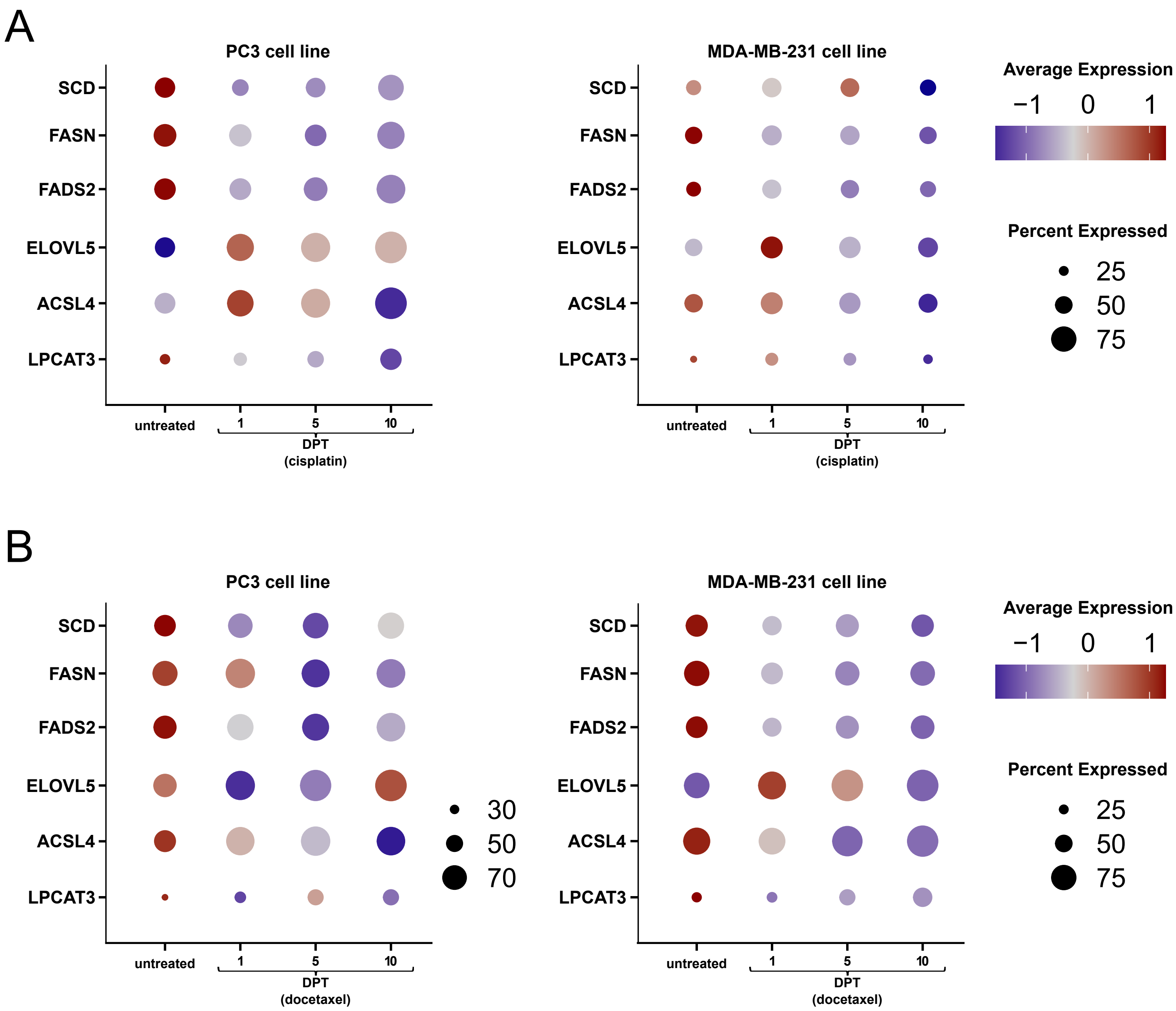


**Supplementary Figure S13. Lipid metabolism enzymes gene expression in cells surviving chemotherapy. A.** Gene expression of ZEB1 driven lipid metabolism enzymes in PC3 or MDA-MB-231 cells surviving cisplatin. Each gene represented by z-score distribution across samples (color) and percentage of cells with detected transcript (size). **B.** Gene expression of ZEB1 driven lipid metabolism enzymes in PC3 or MDA-MB-231 cells surviving docetaxel. Each gene represented by z-score distribution across samples (color) and percentage of cells with detected transcript (size).
Stearoyl-CoA Desaturase (SCD). Fatty Acid Synthase (FASN). Fatty Acid Desaturase 2 (FADS2). ELOVL Fatty Acid Elongase 5 (ELOVL5). Acyl-CoA Synthetase Long Chain Family Member 4 (ACSL4). Lysophosphatidylcholine Acyltransferase 3 (LPCAT3).
